# The flavohemoglobin Yhb1 is a new interacting partner of the heme transporter Str3

**DOI:** 10.1101/2024.03.03.583214

**Authors:** Florie Lo Ying Ping, Tobias Vahsen, Ariane Brault, Raphaël Néré, Simon Labbé

## Abstract

Nitric oxide (NO) is a gaseous molecule that induces nitrosative stress, which can jeopardize cell viability. Yeasts have evolved diverse detoxification mechanisms to effectively counteract NO-mediated toxicity. One mechanism relies on the flavohemoglobin Yhb1, whereas a second one requires the S-nitrosoglutathione reductase Fmd2. However, in this study, we eliminate any contribution from Fmd2 by using strains deleted for this gene. To investigate heme-dependent activation of Yhb1 in response to NO, we use *hem1Δ*-derivative *Schizosaccharomyces pombe* strains lacking the initial enzyme in heme biosynthesis, forcing cells to assimilate heme from external sources. Under these conditions, *yhb1^+^* mRNA levels are repressed in the presence of iron through a mechanism involving the GATA-type transcriptional repressor Fep1. In contrast, when iron levels are low, the transcription of *yhb1^+^*is derepressed and further induced in the presence of the NO donor DETANONOate. Cells lacking Yhb1 or expressing inactive forms of Yhb1 fail to grow in a hemin-dependent manner when exposed to DETANONOate. Similarly, loss of function of the heme transporter Str3 phenocopies the effects of Yhb1 disruption by causing hypersensitivity to DETANONOate under hemin-dependent culture conditions. Coimmunoprecipitation and bimolecular fluorescence complementation assays demonstrate the interaction between Yhb1 and the heme transporter Str3. Collectively, our findings unveil a novel pathway for activating Yhb1, fortifying yeast cells against nitrosative stress.

## Introduction

Heme is a prosthetic group consisting of a protoporphyrin ring, with one iron atom coordinated at its center. The redox-active nature of iron, with its ability to lose (Fe^3+^) and accept one electron (Fe^2+^), makes heme a critical cofactor for a wide variety of enzymes, including catalases and flavohemoglobins (flavoHbs)(Donegan et al. 2019; Dutt et al. 2022; Reddi and Hamza 2016; Severance and Hamza 2009; Swenson et al. 2020). These enzymes play a vital role in protecting cells against harmful reactive oxygen species (ROS) and reactive species of nitrogen (RNS)(Bonamore and Boffi 2008; Glorieux and Calderon 2017). Moreover, heme acts as a small signaling molecule in various processes, including the regulation of the circadian clock, activation of diverse sensor proteins, and gas sensing (Donegan et al. 2019; Mense and Zhang 2006).

On the other hand, maintaining heme homeostasis is essential due to its intrinsic peroxidase activity and its capacity to generate ROS through its redox-active native nature. Moreover, heme can induce cellular damage if it becomes mis-incorporated into proteins or lipid bilayers (Kumar and Bandyopadhyay 2005; Severance and Hamza 2009). Consequently, organisms have developed various mechanisms to safely acquire and distribute heme intracellularly, ensuring appropriate cellular levels and mitigating its potential harmful effects. For instance, in the case of heme prototrophs, these organisms employ a heme biosynthetic pathway that involves eight highly conserved anabolic enzymes(Severance and Hamza 2009). These enzymes are located either in mitochondria or the cytosol, depending on where their specific activities occur within the pathway. A second strategy used by a number of heme auxotrophic organisms is to acquire heme from external sources (Chambers et al. 2021). In contrast to the well-characterized enzymes responsible for endogenous heme synthesis, knowledge of the proteins involved in acquiring exogenous heme is limited and has only been investigated in a small number of organisms.

In the case of the fission yeast *Schizosaccharomyces pombe*, both heme biosynthesis and the acquisition of heme from external sources are pathways through which cells maintain heme homeostasis. To dissociate these processes, a genetic system has been developed in *S. pombe* by creating strains with a deletion of the *hem1^+^* gene (*hem1Δ*), which blocks *de novo* heme biosynthesis by removing the first enzyme, Hem1 (δ-aminolevulinic acid synthase), from the heme biosynthetic pathway (Labbé et al. 2020). With endogenous heme production blocked in these *hem1Δ* cells, two strategies ensure their viability. First, *hem1Δ* cells can be maintained alive by supplementing them with exogenous δ-aminolevulinate (ALA). Assimilation of exogenous ALA enables heme biosynthesis to resume at the second enzymatic step and progress through subsequent enzymatic reactions needed for complete heme biosynthesis. Second, *S. pombe* cells lacking Hem1 (*hem1Δ*) can be kept alive on an ALA-free medium supplemented with exogenous hemin (heme chloride). Under these conditions, *hem1Δ* cells are compelled to rely solely on their own heme uptake machinery.

Using the *hem1Δ*-based genetic system in the absence of ALA and in the presence of hemin, studies have unveiled the existence of two independent heme uptake systems in *S. pombe* (Mourer et al. 2015; Mourer et al. 2017; Normant et al. 2018). One system involves the cell-surface GPI-anchored protein Shu1 (Mourer et al. 2015). When Shu1-expressing *hem1Δ* cells are cultured in the presence of hemin or its fluorescent analog zinc mesoporphyrin IX (ZnMP), Shu1 binds hemin or ZnMP, and undergoes rapid internalization from the cell surface to the vacuolar membrane. Microscopic studies have shown that Shu1 primarily delivers ZnMP to the vacuole prior carrying it further into the cytoplasm in the presence of the vacuolar transporter Abc3 (Mourer et al. 2017). The hemin-mediated vacuolar targeting of Shu1 depends on Ubi4-dependent ubiquitination, the E2 ubiquitin-conjugating enzyme Ubc13, the receptor Nbr1 and the ESCRT (endosomal sorting complex required for transport)-dependent endosomal sorting system, which includes proteins Hse1 and Sst6 (Mourer et al. 2019).

A second heme uptake system relies on the cell-surface transmembrane protein Str3, which belongs to the major facilitator superfamily (MFS) of transporters (Normant et al. 2018). Str3 is predicted to contain 12 transmembrane spans: the first 6 spans form the first half of the transporter, while the remaining 6 spans constitute the second half of the protein. Studies have shown that Str3-mediated transport of exogenous hemin necessitates twice the concentration of hemin (0.15 µM) required by Shu1 (0.075 µM) to rescue the heme-dependent growth deficit of heme auxotrophic cells (Normant et al. 2018). Str3 possesses an atypical extended extracellular loop 11 containing two heme-binding motifs. The amino acid composition of these motifs resembles the near-iron transporter (NEAT) motifs found in several prokaryotic hemeproteins (Honsa et al. 2014). This loop is predicted to serve as an initial heme docking site, facilitating heme passage across the plasma membrane and its entry into the cell. Studies of hemin acquisition in *hem1Δ shu1Δ* cells expressing Str3-GFP have indicated that the plasma membrane localization of Str3-GFP remains unchanged at the cell surface when cells are treated with different concentrations of hemin (Normant et al. 2021). This observation suggests that Str3 remains stable upon exposure to increased concentrations of hemin, implying the involvement of cellular factors in heme mobilization during or after its transport across the plasma membrane by Str3.

Nitric oxide (NO) is a radical gas molecule that can exert either beneficial or detrimental effects within cells, depending on its concentration(Lundberg and Weitzberg 2022). At high concentrations, NO becomes cytotoxic (Tillmann et al. 2011). The fission yeast *S. pombe* can withstand cytotoxic attacks from nitric oxide (NO) through the activity of two enzymes: the flavohemoglobin (flavoHb) Yhb1 and the S-nitrosoglutathione reductase Fmd2 (Astuti et al. 2016). Yhb1 detoxifies NO to nitrate through its NO dioxygenase (NOD) activity. The functional monomeric Yhb1 contains a single N-terminal heme-binding domain followed by successive C-terminal FAD- and NAD(P)-binding domains. In the case of Fmd2, the enzyme reduces S-nitrosoglutathione (GSNO) to ammonia and glutathione disulphide (GSSG) (Liu et al. 2001). Since GSNO can serve as a potential source of bioavailable NO during its decomposition by photolysis or catalysis by reactive metal ions (Lozinsky et al. 2013), Fmd2 prevents the formation of NO from GSNO through its specific GSNO-metabolizing activity. Examination of expression profiles of *yhb1^+^* and *fmd2^+^* genes in *S. pombe* revealed that *yhb1^+^*transcripts are primarily detected during the logarithmic phase and to a lesser extent into the stationary phase. In contrast, *fmd2^+^* mRNA levels are prominently detected after the entry into the stationary phase (Astuti et al. 2016). For both *yhb1^+^* and *fmd2^+^*, mRNA and protein levels transiently increase in the presence of the NO donor DETANONOate (Astuti et al. 2016). Currently, the identity of a major NO-responsive transcriptional regulator in *S. pombe* remains unclear.

Our previous proximity-dependent biotinylation studies on *S. pombe hem1Δ shu1Δ str3Δ* cells expressing a functional *str3^+^-BirA* allele have uncovered an interaction between Yhb1 and Str3 in the presence of hemin under low iron conditions (Normant et al. 2021). Although heme is distributed to heme-requiring enzymes like Yhb1 upon uptake into cells, the specific mechanisms by which cellular hemeproteins acquire exogenous heme remain unknown.

In the present study, we determined that mRNA levels of *yhb1^+^* are increased under low iron conditions, whereas *yhb1^+^* transcripts exhibit reduced expression levels under both basal and iron-replete conditions. Moreover, our findings showed that the transcriptional repressor Fep1 plays a role in the iron-dependent regulation of the *yhb1^+^* gene. Under conditions of hemin-dependent growth, Yhb1 is required for counteracting the growth inhibition of *S. pombe hem1Δ fmd2Δ* cells induced by DETANONOate. Coimmunoprecipitation and bimolecular fluorescence complementation assays revealed that Yhb1 interacts with Str3 when cells utilize hemin as the sole source of heme under low iron conditions. Collectively, these results unveil a route through which heme is delivered to Yhb1 via its interaction with the heme transporter Str3 at the plasma membrane in response to nitrosative stress under conditions of low iron and hemin dependence.

## Results

### The expression of yhb1^+^ is downregulated in response to iron in cells deficient in heme biosynthesis

In the yeast *S. pombe*, transcriptomic analyses have shown that several genes encoding heme-binding proteins are repressed under conditions of excess iron (Mercier et al. 2008; Rustici et al. 2007). Given that the *yhb1^+^* gene encodes for a hemeprotein, we assessed whether *yhb1^+^* expression was differentially regulated as a function of iron availability. To conduct these experiments, we used cells lacking the *hem1^+^* gene (*hem1Δ*) in which production of endogenous heme is blocked, requiring cells to rely on exogenous hemin as their sole source of heme for cell growth. Under these conditions, *hem1Δ* cells were either left untreated or exposed to the iron chelator 2,2’-dipyridyl (Dip, 250 µM) or FeCl_3_ (Fe, 250 µM) for 3 h. In the presence of Dip, *yhb1^+^* mRNA levels were induced 3.2-fold compared to basal levels of expression in untreated cells (Fig. 1, A – B). In the presence of iron, *yhb1^+^* mRNA levels were similar to those observed in untreated cells (Fig. 1, A – B).

**Fig. 1.**
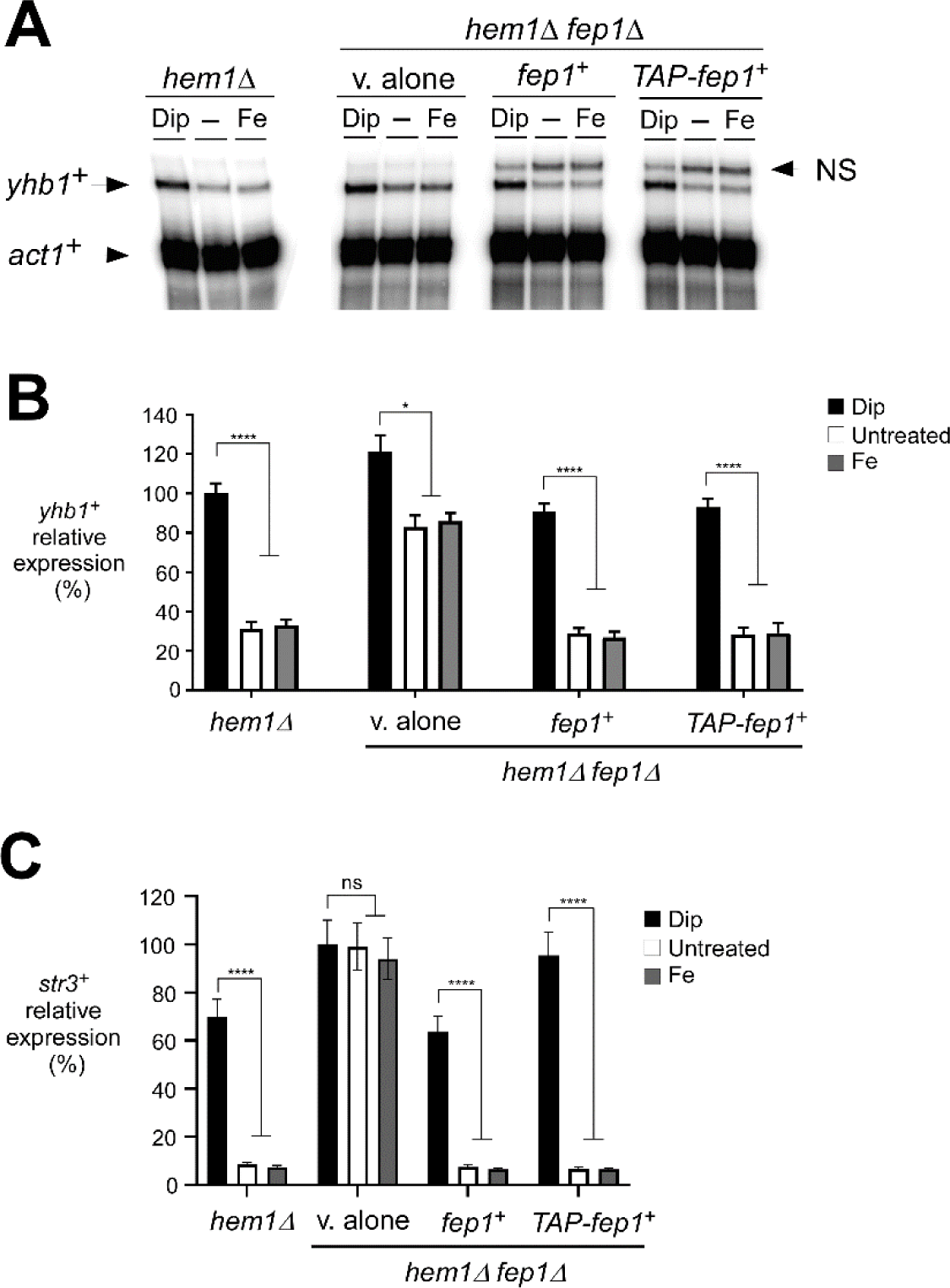
*The iron-responsive regulator Fep1 plays a role in repressing the expression of yhb1^+^ in cells incapable of synthesizing heme*. *A*, The indicated yeast strains were transferred in ALA-free medium for 5 h. Following washes, cultures were incubated with hemin (5 µM) without further supplementation (-), or with Dip (250 µM) or FeCl_3_ (Fe, 100 µM) for 3 h. Total RNA was extracted from culture aliquots, and steady-state mRNA levels of *yhb1^+^* and *act1^+^*were analyzed by RNase protection assays. “NS”, indicates nonspecific signal. *B*, The graph depicts the results quantified from three independent RNase protection assays. Values are presented as averages ± S.D. Asterisks denote statistical significance, with ** indicating p < 0.01, and **** indicating p < 0.0001 (determined by 1-way ANOVA with Dunnett’s multiple comparisons test against the Dip-treated strains). *C*, Aliquots of RNA samples were used for the assessment of *str3^+^* mRNA levels using RT-qPCR assays. *str3^+^* served as a control gene known to be regulated by iron and controlled by Fep1. The graph depicts the quantification results from three separate RT-qPCR experiments. Values are presented as described in *panel B*. “NS”, stands for no statistical significance.

To further investigate whether Fep1 played a role in regulating the *yhb1^+^* gene, we examined the impact of disrupting *fep1^+^* (*fep1Δ*) on *yhb1^+^* expression when *hem1Δ* cells were cultured with exogenous hemin as their exclusive source of heme. For this purpose, *hem1Δ fep1Δ* mutant cells were transformed with a control integrative plasmid (v. alone) or a plasmid harboring an untagged or a TAP-tagged version of Fep1 under the control of its own promoter. These cells were subjected to the same conditions and treatments as described in the case of *hem1Δ* cells. After 3 h, total RNA was isolated and we analyzed *yhb1^+^* gene expression using RNase protection assays. The loss of Fep1 led to a notable increase in the levels of *yhb1^+^*expression, particularly under both basal and iron-replete conditions (Fig. 1, A – B). Although the derepression of *yhb1^+^* was not as pronounced as in iron-starved conditions (Dip), it was higher than the expression observed under the same basal and iron-replete conditions in *hem1Δ* cells expressing endogenous *fep1^+^* (2.7-fold) or in *hem1Δ fep1Δ* cells where functional *fep1^+^* and *TAP-fep1^+^* alleles were reintegrated (2.9-fold) (Fig. 1, A – B).

To serve as a control, we concurrently analyzed *str3^+^* transcripts. These transcripts are known to be induced during iron starvation conditions and repressed when iron levels are high, a process regulated by Fep1 activity. As expected, *str3^+^* mRNA levels were repressed under high-iron conditions compared to their corresponding levels under low-iron conditions in *hem1Δ* cells expressing the endogenous *fep1^+^* (9.5-fold less) or in *hem1Δ fep1Δ* cells expressing untagged and TAP-tagged *fep1^+^* (10.0- and 14.9-fold less, respectively) (Fig. 1C). In contrast, when analyzed in *hem1Δ* cells lacking a functional *fep1^+^* allele (*fep1Δ*), *str3^+^*mRNA levels remained high and relatively unchanged under all experimental conditions (Fig. 1C). Taken together, these results showed that *yhb1^+^* is a gene subject to iron regulation, with Fep1 exerting an iron-dependent inhibitory effect on Yhb1 expression.

### Fep1 occupies the yhb1^+^ promoter in response to high concentrations of iron

To further investigate whether Fep1 directly represses *yhb1^+^*expression through its recruitment to the *yhb1^+^* promoter, we employed a *hem1Δ php4Δ fep1Δ* mutant strain that expressed both untagged and TAP-tagged *fep1^+^* alleles. It is worth noting that this mutant strain lacks Php4, resulting in the constitutive expression of reintegrated *fep1^+^* alleles (Mercier and Labbe 2009; Mercier et al. 2008). This approach ensures that any observed effects of iron on Fep1 or TAP-Fep1 remain unaffected by potential changes in the transcription of the *fep1^+^* or *TAP-fep1^+^*gene, as it occurs independently of Php4 (Mercier et al. 2008). Using this biological system, we examined whether there was presence of TAP-Fep1 in the promoter region of *yhb1^+^* using a ChIP method (Fig. 2, A – B). The ChIP analysis revealed that when cells were cultured in the presence of hemin (5 µM) and treated with FeCl_3_ (100 µM) for 3 h, TAP-Fep1 immunoprecipitated chromatin corresponding to the *yhb1^+^* promoter 6.8-fold more than an 18S ribosomal region reference (Fig. 2B). In contrast, when *hem1Δ php4Δ fep1Δ* cells expressing TAP-Fep1 were incubated in the presence of Dip, only weak levels of *yhb1^+^*promoter fragments were immunoprecipitated (Fig. 2B). Importantly, the amount of immunoprecipitated chromatin by TAP-Fep1 at the *yhb1^+^* promoter was higher when cells were treated with iron (10.0-fold increase) compared to Dip (Fig. 2B). The occupation of the *yhb1^+^* promoter by TAP-Fep1 was detected using primers that amplify a DNA region located between positions -1486 and -1385 relative to the *yhb1^+^* initiator codon (Fig. 2A). This promoter region contains predicted GATA elements that are known to serve as potential binding sites for Fep1.

**Fig. 2.**
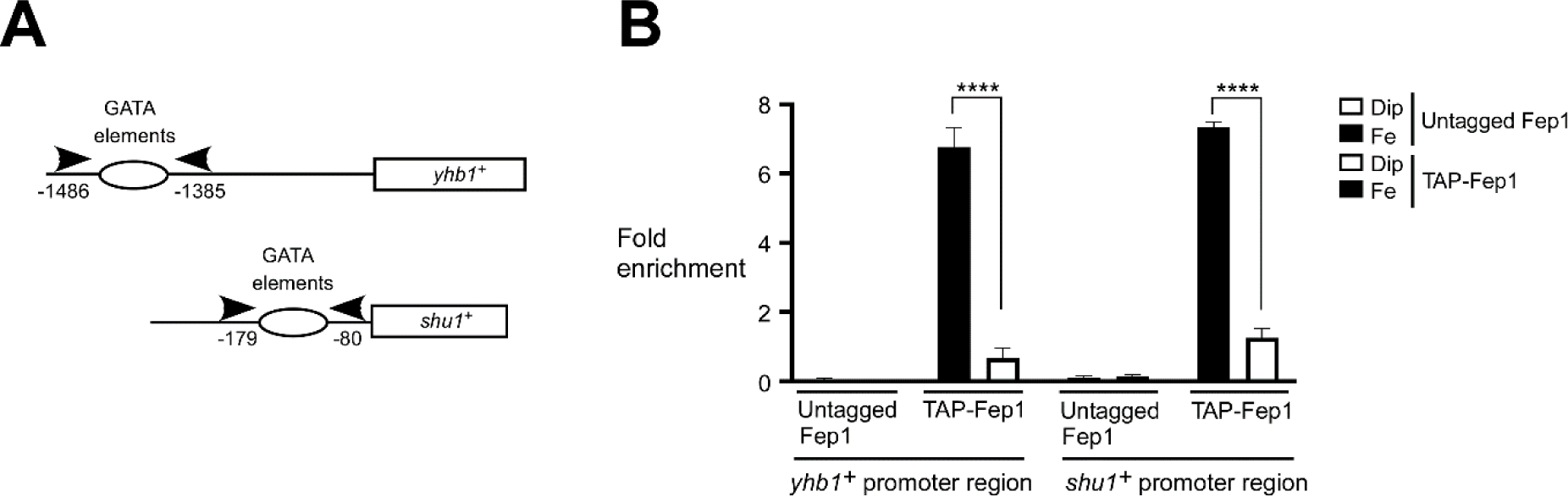
*Fep1 interacts with the yhb1^+^ promoter under iron-replete conditions*. *A*, Schematic representation of the *yhb1^+^* and *shu1^+^* promoter regions. Arrows indicate primer locations for qPCR analysis. Nucleotide numbers refer to the position relative to the A of the initiator codon of each gene (*yhb1^+^* or *shu1^+^*). Empty ovals indicate promoter regions containing GATA elements known to serve as binding sites for Fep1. *B*, *hem1Δ php4Δ fep1Δ* cells expressing an integrated untagged or TAP-tagged *fep1^+^* allele were transferred to an ALA-free medium containing FeCl_3_ (75 µM) for 5 h. After washing, the cultures were incubated in YES medium supplemented with hemin (5 µM) and treated with Dip (250 µM) or FeCl_3_ (Fe, 100 µM) for 3 h. Subsequently, cultures were fixed by formaldehyde treatment. Cell lysates were prepared by glass bead disruption and chromatin was prepared and immunoprecipitated using Sepharose-bound anti-mouse IgG antibodies. Binding of TAP-Fep1 to the *yhb1^+^* and *shu1^+^* promoters was calculated by measuring the enrichment of specific *yhb1^+^* (positions -1486 to -1385) and *shu1^+^* (positions -179 to -80) regulatory promoter regions relative to an 18S ribosomal DNA coding region. ChIP data were calculated as values of the largest amount of chromatin measured (fold enrichment). Results are shown as averages ± SD from a minimum of three independent experiments, each performed in biological triplicate. Asterisks indicate statistical significance (****, p < 0.0001, one-way ANOVA with Dunnett’s multiple comparisons test against iron-replete cells expressing TAP-Fep1).

As positive control experiments, the ChIP analysis demonstrated that in the case of *hem1Δ php4Δ fep1Δ* cells expressing TAP-Fep1 and treated with FeCl_3_ (100 µM), TAP-Fep1 occupied the *shu1^+^*promoter (a known target of Fep1) with a 7.3-fold enrichment relative to a control 18S ribosomal DNA region reference (Fig. 2B). In contrast, the association of TAP-Fep1 with the *shu1^+^* promoter was much weaker (1.2-fold) under low-iron conditions compared to an 18S ribosomal region reference (Fig. 2B). This lower enrichment level was 5.9-fold less than what was observed in cells expressing TAP-Fep1 and incubated with iron (Fig. 2B). As negative controls, untagged Fep1 immunoprecipitated only background levels of the *yhb1^+^* and *shu1^+^* promoter regions (Fig. 2B). Taken together, these results showed that the *yhb1^+^* promoter is bound by Fep1 under iron-replete conditions and in the presence of hemin as the sole source of heme.

### Effect of DETANONOate on the expression of yhb1^+^

Previous studies had established that *S. pombe* can counter NO cytotoxic attacks, primarily through the activity of two enzymes: the NO dioxygenase Yhb1 and the S-nitrosoglutathione reductase Fmd2 (Astuti et al. 2016). Given the overlapping functions of these enzymes, we generated mutant strains to eliminate the potential interference from Fmd2. This was achieved by systematically inactivating the *fmd2^+^* allele (*fmd2Δ*) in the *hem1Δ*-derived strains used for subsequent analyses. With the goal of further characterizing Yhb1 during hemin-dependent cell growth, we examined whether exposure to NO led to the transcriptional activation of *yhb1^+^* under these growth conditions. *hem1Δ fmd2Δ* or *hem1Δ yhb1Δ fmd2Δ* cells expressing untagged and TAP-tagged *yhb1^+^* alleles were precultured and then transferred to ALA-free medium containing hemin (1 µM) until reaching an OD_600_ of 0.8. At this stage, the cultures were shifted to hemin-supplemented medium with a pH adjusted to 7.2, and subjected to DETANONOate treatment for 0, 60, 90, and 180 min. Aliquots of cultures were collected and total RNA isolated at the indicated time points. The expression of *yhb1^+^* was assessed by RT-qPCR assays. Results showed that in DETANONOate-treated *hem1Δ fmd2Δ* cells, the expression of *yhb1^+^* was induced 2.4-, 2.7-, and 3.4-fold above the levels observed in untreated cells after 60, 90, and 180 min, respectively (Fig. 3A). A similar pattern of DETANONOate-induced expression was observed for both *yhb1^+^* (2.4-, 2.7-, and 3.5-fold increase) and *yhb1^+^-TAP* (2.5-, 2.7-, and 3.7-fold increase) transcripts when probed in *hem1Δ yhb1Δ fmd2Δ* cells under the same experimental conditions (Fig. 3A).

**Fig. 3.**
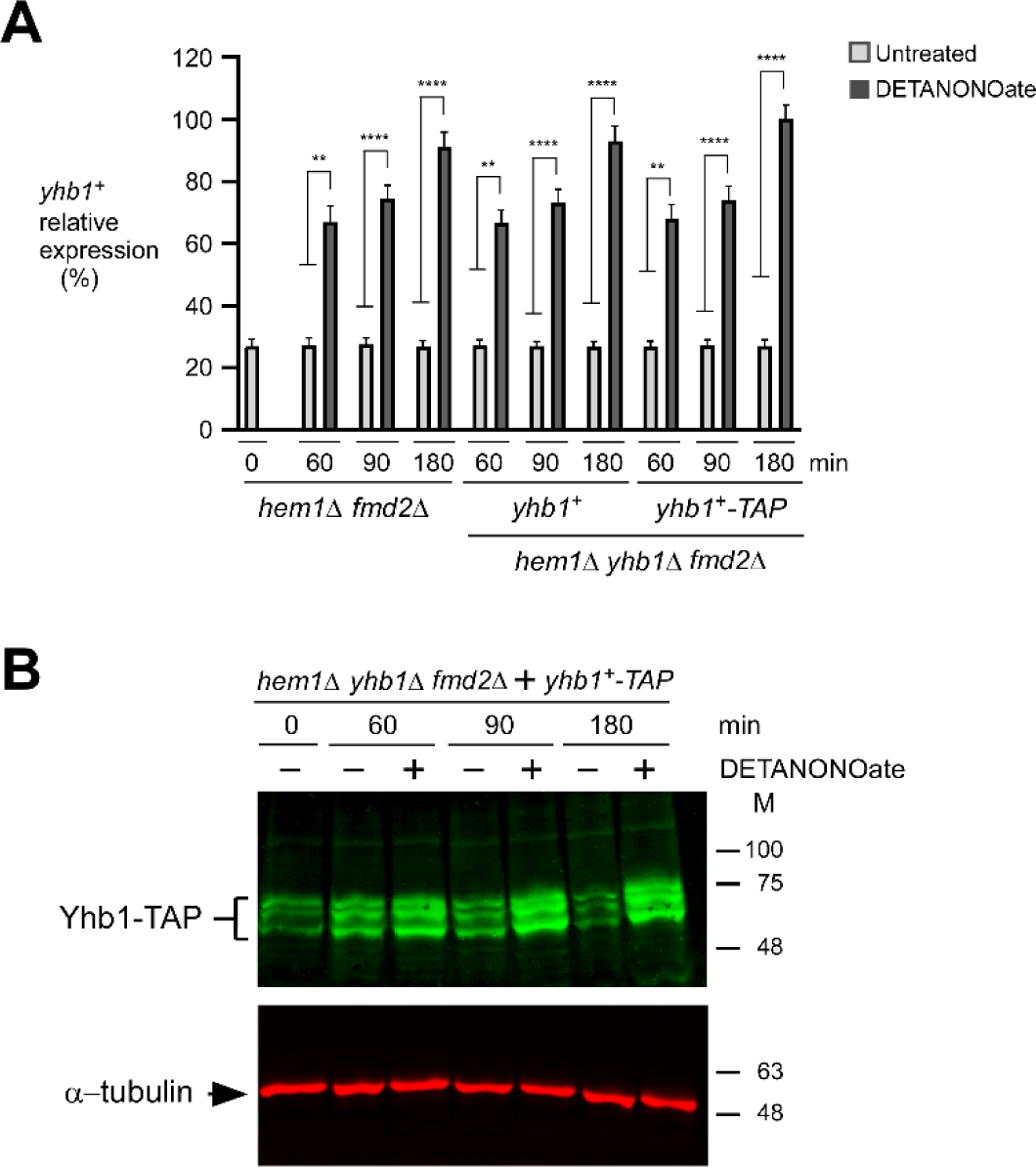
*Yhb1 steady-state mRNA and protein levels are induced upon exposure to exogenous DETA NONOate under conditions of low iron*. *A*, *hem1Δ fmd2Δ* and *hem1Δ yhb1Δ fmd2Δ* strains expressing the specified alleles were cultured in ALA-free medium supplemented with Dip (50 µM) and hemin (1 µM) for 6 h. After washes, the cultures were transferred to YES medium adjusted to pH 7.2 and then treated with DETA NONOate (3 mM) in the presence of Dip (50 µM) and hemin (1 µM). The zero time point (0 min) refers to onset of DETA NONOate treatment. Total RNA was isolated from culture aliquots collected at the indicated time points. Following RNA extraction, steady-state mRNA levels of *yhb1^+^*, *yhb1^+^-TAP*, and *act1^+^* were assessed through RT-qPCR assays. The graph presents quantification data from three independent RT-qPCR experiments, with error bars indicating standard deviations (± SD). Asterisks correspond to p < 0.01 (**) and p < 0.0001 (****) (determined by two-way ANOVA with Tuckey’s multiple comparisons test, comparing to the indicated strain treated with DETA NONOate). *B*, In the case of *hem1Δ yhb1Δ fmd2Δ* cells expressing the *yhb1^+^-TAP* allele, whole cell extracts from culture aliquots used in *panel A* were subjected to immunoblot assays. These assays utilized a polyclonal anti-mouse IgG antibody (recognizing Yhb1-TAP) and a monoclonal anti-α-tubulin antibody. Molecular weight standards (M) are indicated to the right of the panel.

To investigate whether the steady-state protein levels of Yhb1-TAP followed those of *yhb1^+^* mRNA upon exposure to DETANONOate, we used *hem1Δ yhb1Δ fmd2Δ* cells, in which *yhb1^+^-TAP* was reintegrated. Results showed that the steady-state protein levels of Yhb1-TAP increased after 60, 90, and 180 min, exhibiting 1.4-, 2.9-, and 3.8-fold higher enrichment compared to untreated cells (Fig. 3B). These findings collectively indicated that DETANONOate induces the expression of *yhb1^+^*, subsequently elevating the steady-state levels of its protein product in response to nitrosative stress.

### Yhb1 protects hem1Δ yhb1Δ fmd2Δ mutant cells from nitrosative stress

Previous studies have demonstrated that fungal Yhb1-like flavohemoglobins effectively convert NO into harmless nitrate through their heme-dependent dioxygenase activity(de Jesús-Berríos et al. 2003; Liu et al. 2000; Ullmann et al. 2004). Considering their dependency on heme, our study centered on whether Yhb1 remains active when cells solely rely on external sources of heme for their growth. In *S. pombe*, heme comes from two distinct sources: heme biosynthesis and the assimilation of exogenous hemin from the environment. To dissociate these processes, we used *S. pombe* strains with a disruption in the *hem1^+^* gene, responsible for encoding the initial enzyme in the heme biosynthesis pathway. The absence of Hem1 (*hem1Δ*) effectively blocks *de novo* heme production. To compensate this deficiency, we added exogenous hemin into the growth medium. Under these experimental conditions (*hem1Δ* + hemin), *hem1Δ* cells can assimilate exogenous hemin through two independent heme uptake systems requiring Shu1 and Str3, respectively (Labbé et al. 2020). We therefore examined whether heme synthesis-deficient strains expressing a functional *yhb1^+^* allele could grow in the presence of NO when exogenous hemin served as their sole source of heme.

As part of initial controls, we confirmed that a *hem1Δ* strain, when cultured in an ALA-free medium without hemin supplementation, displayed no growth in contrast to *hem1Δ* cells supplemented with either exogenous ALA or hemin (Fig. 4, A and B). Additionally, we validated that other *hem1Δ*-derived strains harboring additional gene disruptions, such as *hem1Δ yhb1Δ*, *hem1Δ fmd2Δ*, *hem1Δ yhb1Δ fmd2Δ*, and *hem1Δ yhb1Δ fmd2Δ* expressing an untagged *yhb1^+^* allele, exhibited comparable growth patterns to *hem1Δ* cells in the presence of exogenous ALA or hemin (Fig. 4, A and B).

**Fig. 4.**
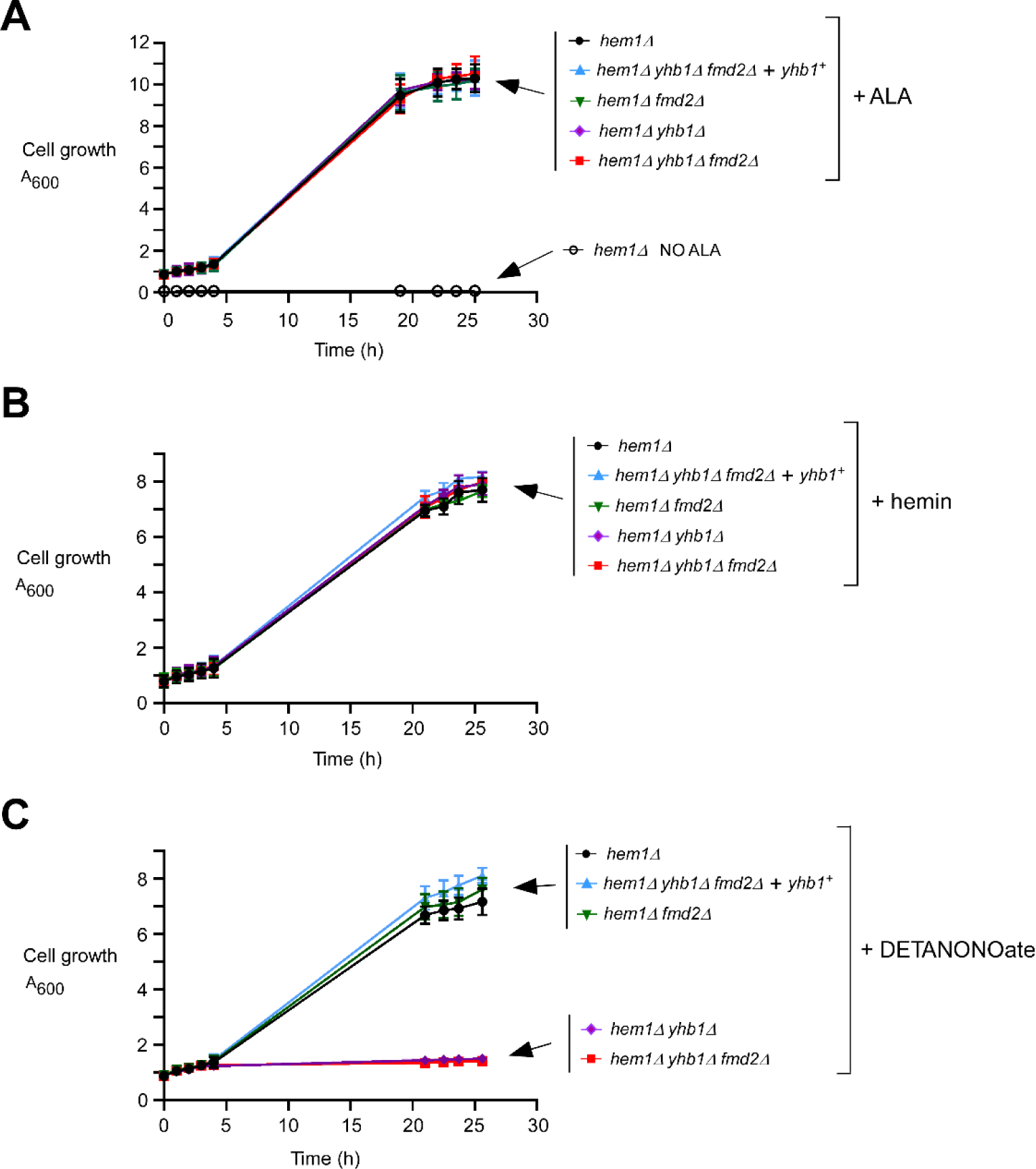
*hem1Δ yhb1Δ fmd2Δ mutant cells require a functional yhb1^+^ allele for growth in the presence of DETANONOate*. *A*, Growth of the indicated yeast strains was assessed in YES medium that was left untreated (no ALA; *open circles*) or supplemented with exogenous ALA (200 µM)*. B – C*, Cell proliferation assays were performed using the indicated strains grown in ALA-free medium supplemented with hemin (1 µM). Once the cultures reached an A_600_ of 0.8, they were transferred to YES medium (pH 7.2) containing hemin (1 µM) without DETA-NONOate supplementation (*panel B*) or with DETANONOate supplementation (3 mM) (*panel C*). Growth of the strains was monitored at the specified time points. Strain color codes are as follows: black, *hem1Δ*; red, *hem1Δ yhb1Δ fmd2Δ*; blue, *hem1Δ yhb1Δ fmd2Δ* expressing *yhb1^+^*; green, *hem1Δ fmd2Δ*; violet, *hem1Δ yhb1Δ*. The graphs represent the results of three independent experiments performed in biological triplicate, with values presented as averages ± SD.

We next tested whether the above-mentioned strains derived from *hem1Δ* were capable of growth when exposed to DETANONOate (3 mM) in an ALA-free medium supplemented with exogenous hemin (1 µM). Results indicated that both *hem1Δ* and *hem1Δ fmd2Δ* strains, harboring an endogenous *yhb1^+^* allele, exhibited cell growth up to an A_600_ of 6.8 and 7.3 after 25.6 h (Fig. 4C). In the case of a *hem1Δ yhb1Δ fmd2Δ* strain expressing an untagged *yhb1^+^* allele, it exhibited similar growth to the positive control, the *hem1Δ* strain (reaching an A_600_ of 7.9 after 25.6 h) (Fig. 4C). In contrast, both the *hem1Δ yhb1Δ* and *hem1Δ yhb1Δ fmd2Δ* strains displayed poor growth (A_600_ of 1.4 after 25.6 h) compared to the *hem1Δ* strain possessing the endogenous *yhb1^+^* gene (Fig. 4C). Collectively, these results indicated that Yhb1 is required for enabling the growth of heme synthesis-deficient cells when supplemented with exogenous hemin in the presence of the nitric oxide donor DETANONOate.

### In the presence of the non-iron metalloporphyrin ZnMP, hem1Δ yhb1Δ fmd2Δ cells expressing Yhb1-mNeonGreen exhibit hypersensitivity to DETANONOate

To further investigate the function of Yhb1, we created different epitope-tagged Yhb1 fusion proteins. To verify that the insertion of these epitopes did not interfere with Yhb1 function, we performed cell proliferation assays. These assays were carried out using a *hem1Δ yhb1Δ fmd2Δ* mutant strain in which untagged *yhb1^+^* and *TAP*-, *Cherry*-, and *mNeonGreen*-tagged *yhb1^+^* alleles were introduced by integration. As control conditions, we performed growth assays with all the indicated strains in ALA-free medium with hemin supplementation (1 µM) (Fig. 5A). Results showed that all tested strains carrying a disrupted *hem1Δ* allele exhibited a robust hemin-dependent growth to an A_600_ of 7.8 to 8.0 after 25 h (Fig. 5A). We next used the same group of strains that were cultured under the same conditions, but in the presence of DETANONOate (3 mM) (Fig. 5B). Prior to treatment, strains were grown to an A_600_ of 0.75 in YES lacking ALA with hemin supplementation. At this 0 h time point (A_600_ of 0.75), strains were treated with DETANONOate or were left without treatment to compare their relative growth (Fig. 5, A – B). In the case of *hem1Δ yhb1Δ fmd2Δ* cells, they exhibited poor growth in a medium supplemented with 3 mM DETANONOate. These cells displayed 4.1-, 4.3-, and 4.2-fold less growth after 19, 23.5, and 25 h, respectively, compared with untreated *hem1Δ yhb1Δ fmd2Δ* cells (Fig. 5, A – B). In contrast, the growth defect observed in the case of DETANONOate-treated *hem1Δ yhb1Δ fmd2Δ* cells was reversed by expression of integrated untagged *yhb1^+^*and *TAP*-, *Cherry*-, or *mNeonGreen*-tagged *yhb1^+^*alleles in a manner comparable to a reference *hem1Δ* strain expressing an endogenous *yhb1^+^*allele (A_600_ of 7.6 to 7.8 after 25 h) (Fig. 5B).

**Fig. 5.**
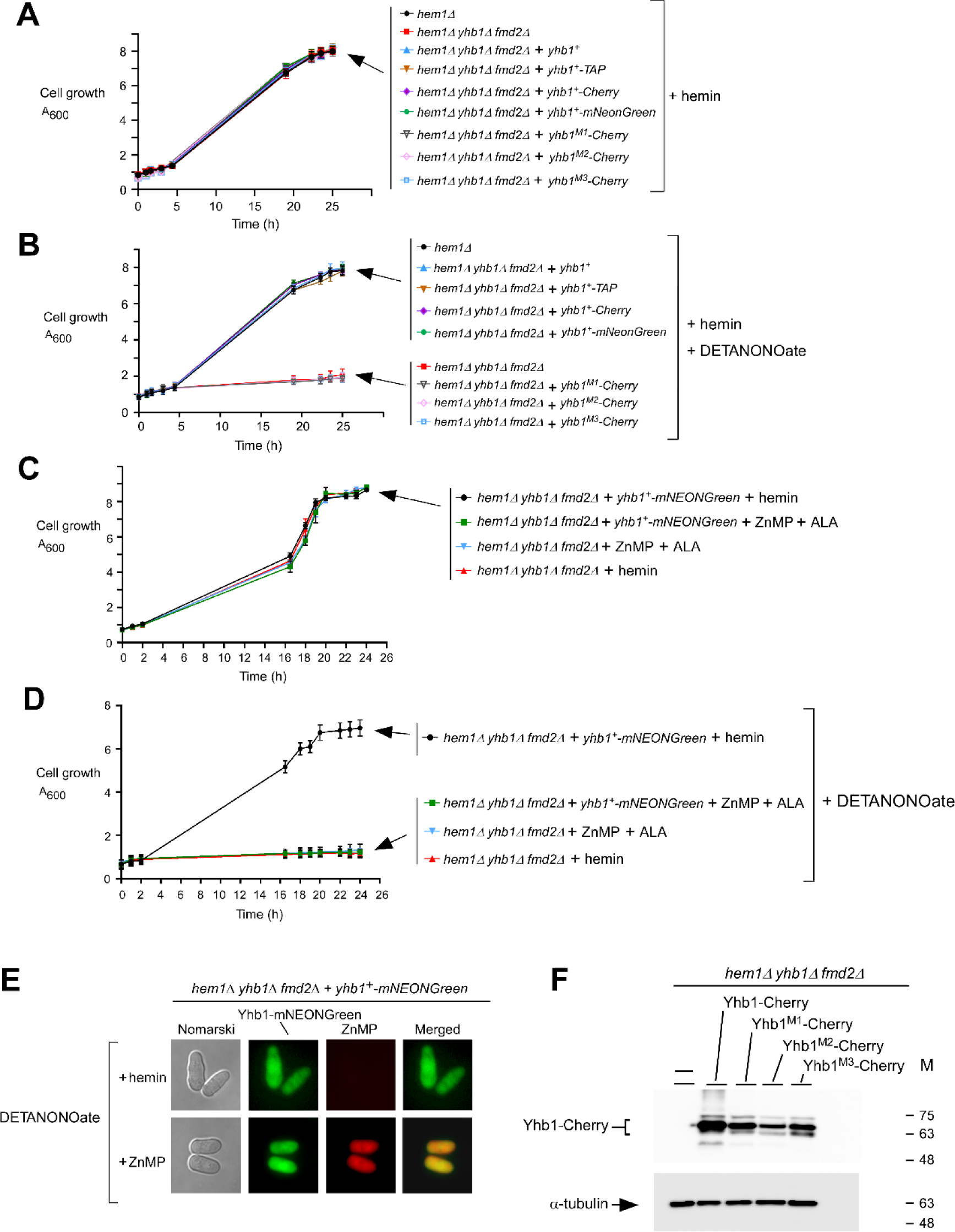
*Growth inhibition of hem1Δ yhb1Δ fmd2Δ cells expressing Yhb1-mNEONGreen upon exposure to ZnMP and DETA-NONOate*. *A – B*, The indicated cell cultures were grown to logarithmic phase (A_600_ of 0.8) in ALA-free medium supplemented with hemin (1 µM). Subsequently, these cultures were transferred to YES medium (pH 7.2) containing hemin (1 µM) without DETANONOate supplementation (*panel A*) or with DETANONOate supplementation (3 mM) (*panel B*). The growth of the specified strains was monitored at the indicated time points. Strain color codes are as follows: black, *hem1Δ*; red, *hem1Δ yhb1Δ fmd2Δ*; blue, *hem1Δ yhb1Δ fmd2Δ* expressing *yhb1^+^*; brown, *hem1Δ yhb1Δ fmd2Δ* expressing *yhb1^+^-TAP*; violet, *hem1Δ yhb1Δ fmd2Δ* expressing *yhb1^+^-Cherry*; green, *hem1Δ yhb1Δ fmd2Δ* expressing *yhb1^+^-mNeonGreen*; gray unfilled triangle, *hem1Δ yhb1Δ fmd2Δ* expressing *yhb1^M1^-Cherry*; pink unfilled diamond, *hem1Δ yhb1Δ fmd2Δ* expressing *yhb1^M2^-Cherry*; and, blue unfilled square, *hem1Δ yhb1Δ fmd2Δ* expressing *yhb1^M3^-Cherry*. *C – D*, *hem1Δ yhb1Δ fmd2Δ* cells harboring an empty vector or expressing Yhb1-mNeonGreen were divided into four treatment groups. One group was incubated in the presence of ALA (1 µM) and ZnMP (10 µM) but without DETANONOate treatment. The second group was incubated in the presence of hemin (1 µM) but without DETANONOate treatment (*panel C*). The third group was exposed to ALA (1 µM) along with a combination of ZnMP (10 µM) and DETANONOate (3 mM). The fourth group was treated with hemin (1 µM) and DETANONOate (3 mM) (*panel D*). The growth of the specified strains was monitored at the indicated time points. Strain color codes are as follows: red, *hem1Δ yhb1Δ fmd2Δ* in the presence of hemin; blue, *hem1Δ yhb1Δ fmd2Δ* in the presence of ZnMP and ALA; black, *hem1Δ yhb1Δ fmd2Δ* expressing *yhb1^+^-mNEONGreen* in the presence of hemin; and green, *hem1Δ yhb1Δ fmd2Δ* expressing *yhb1^+^-mNEONGreen* in the presence of ZnMP and ALA. The graph represents the results of three independent experiments, each performed in biological triplicate, with values presented as averages ± SD. *E*, At the 16-h time point, aliquots of the cultures used in *panel D* were subjected to fluorescence microscopy to visualize the cellular localization of Yhb1-mNEONGreen (center left) and the cytoplasmic accumulation of ZnMP (center right) within the cells. Cell morphology was examined using Nomarski optics (far left). Merged images of Yhb1-mNEONGreen and ZnMP fluorescent signals are displayed in the far-right panels. The results of microscopy are representative of five independent experiments. *F*, After 19 h of incubation in an ALA-free medium supplemented with hemin (1 µM) under low-iron conditions, aliquots of *hem1Δ yhb1Δ fmd2Δ* cells expressing the indicated wild-type and yhb1^M1-, M2-, M3-^Cherry mutant alleles were subjected to whole cell extract preparations. Cell lysates were analyzed by immunoblot assays using anti-Cherry and anti-α-tubulin antibodies. Molecular weight standards (in kDa) are indicated on the right side of the panels.

We next used *hem1Δ yhb1Δ fmd2Δ* cells expressing either *yhb1^M1^-Cherry*, *yhb1^M2^-Cherry*, or *yhb1^M3^-Cherry* mutant allele to further validate that wild-type Yhb1 played a role in detoxifying DETANONOate when cells were cultured in an ALA-free medium supplemented with exogenous hemin under low iron conditions. According to previous structural studies performed on bacterial and fungal flavoHbs, several amino acid residues are conserved in *S. pombe* Yhb1 and may serve as potential heme ligands(El Hammi et al. 2012; Ilari and Boffi 2008). To validate their potential role in Yhb1 activity, different combinations of mutated residues were created by substituting them with Ala residues. The residue His^114^ was replaced by an Ala, resulting in the Yhb1^M1^-Cherry mutant. In the case of the Yhb1^M2^-Cherry mutant, amino acid residues Tyr^58^, His^114^, Tyr^124^, Pro^125^, Ile^126^, and Val^127^ were replaced by alanine residues, whereas the Yhb1^M3^-Cherry mutant contained the same substitutions as Yhb1^M2^-Cherry, with an additional substitution where Glu^167^ was replaced with an Ala. When *hem1Δ yhb1Δ fmd2Δ* cells expressing these mutant alleles were treated with DETANONOate, they were hypersensitive to DETANONOate, exhibiting poor growth that was comparable to *hem1Δ yhb1Δ fmd2Δ* cells containing an empty plasmid (Fig. 5B). After 19 h, whole-cell extracts were prepared and analyzed by immunoblotting. The results showed that mutant derivatives of Yhb1 were produced in *hem1Δ yhb1Δ fmd2Δ* cells (Fig. 5F). Taken together, these results showed that a functional untagged or epitope-tagged *yhb1^+^*is required to allow *hem1Δ yhb1Δ fmd2Δ* cells to grow in ALA-free medium supplemented with hemin and DETANONOate.

In *S. pombe*, the non-iron metalloporphyrin zinc mesoporphyrin (ZnMP) is taken up via two heme uptake pathways that require Shu1 and Str3, respectively (Mourer et al. 2015; Normant et al. 2018). High concentrations of ZnMP are cytotoxic to yeast cells, presumably due to its interference with essential hemeproteins (Hu et al. 2013; Roy et al. 2022; Stojiljkovic et al. 2001). Building on this observation, we investigated whether ZnMP assimilation could impede Yhb1 function, rendering yeast cells hypersensitive to DETANONOate (a NO donor).

Cells lacking *hem1^+^*, *yhb1^+^*, and *fmd2^+^* genes (*hem1Δ yhb1Δ fmd2Δ*) harboring an empty plasmid or expressing Yhb1-mNeonGreen were precultured in the presence of ALA until reaching logarithmic phase (A_600_ of ∼0.8). Subsequently, these cells were washed and divided into four treatment groups. One group was incubated in the presence of ALA (1 µM) and ZnMP (10 µM) but without nitrosative stress. The second group was incubated in the presence of hemin (1 µM) but without nitrosative stress. Both of these initial groups exhibited robust growth (Fig. 5C). The third group was exposed to ALA (1 µM) along with a combination of ZnMP (10 µM) and DETANONOate (3 mM). The fourth group was treated with hemin (1 µM) and DETANONOate (3 mM). Notably, after 24 hours, cells supplemented with hemin in the fourth group displayed significant growth, reaching an A_600_ of 7.2 in the presence of DETANONOate (see Fig. 5D). In contrast, cells in the third group treated with ZnMP exhibited a growth defect when exposed to DETANONOate (Fig. 5D). Control experiments using the *hem1Δ yhb1Δ fmd2Δ* triple mutant consistently showed no growth in the presence of DETANONOate, regardless of hemin or ALA plus ZnMP supplementation, confirming the impact of DETANONOate treatment (Fig. 5D).

To confirm the expression of Yhb1-mNeonGreen and the assimilation of ZnMP in *hem1Δ yhb1Δ fmd2Δ* triple mutant cells, we analyzed culture aliquots from the indicated cultures using fluorescence microscopy (Fig. 5E). The results revealed that DETANONOate-treated cells carrying the integrated *yhb1^+^-mNEONGreen* allele exhibited a green fluorescence signal distributed across the cell when exposed to either exogenous hemin or ZnMP (Fig. 5E). Parallel fluorescence microscopy analysis of culture aliquots determined that cells incubated with ZnMP displayed significant accumulation of the fluorescent heme analog within their cytosol (Fig. 5E). These results showed that *hem1Δ yhb1Δ fmd2Δ* cells, following transformation with the mNEONGreen epitope-tagged *yhb1^+^* allele, generate a Yhb1-mNEONGreen-associated fluorescence signal that largely colocalizes with the ZnMP fluorescence detected in the cytosol of cells. Furthermore, use of the non-iron metalloporphyrin ZnMP inactivates Yhb1, rendering Yhb1-mNEONGreen-expressing *hem1Δ yhb1Δ fmd2Δ* cells hypersensitive to NO.

### Subcellular localization of Yhb1 and Str3 under low-iron conditions and in the presence of exogenous hemin

Our prior work using a BioID2 proximity-dependent biotinylation system identified Yhb1 as an interacting partner protein of the heme transporter Str3 (Normant et al. 2021). Based on these findings, we created a *hem1Δ shu1Δ str3Δ yhb1Δ fmd2Δ* mutant strain in which the *yhb1^+^-Cherry* allele was cointegrated with the *str3^+^-GFP* allele. Once these cointegrated cells reached mid-logarithmic phase, we transferred them to an ALA-free medium for a 6 h period. Within the final 3 h of incubation, aliquots of cells were treated with FeCl_3_ (100 µM), Dip (250 µM) or Dip supplemented with hemin (1 µM). For each treatment group, we assessed the cellular localization of Yhb1-Cherry and Str3-GFP. The red fluorescence signal associated with Yhb1-Cherry was primarily observed throughout the cytoplasm and it appeared to be largely absent within vacuoles in the presence of Dip or a combination of Dip and hemin (Fig. 6). Furthermore, a significant decrease in fluorescence was observed upon iron treatment. In the case of the Str3-GFP-associated fluorescence signal, it predominantly localized at the cell periphery when cells were treated with Dip or Dip plus hemin (Fig. 6). In contrast, the green fluorescence signal of Str3-GFP was lost under iron-replete conditions, consistent with the absence of *str3^+^* mRNA expression when cells are exposed to exogenous iron (Normant et al. 2018). Taken together, these findings suggested that a fraction of cytosolic Yhb1 may potentially interact with Str3 at the plasma membrane when iron-starved cells rely on exogenous hemin as their sole source of heme.

**Fig. 6.**
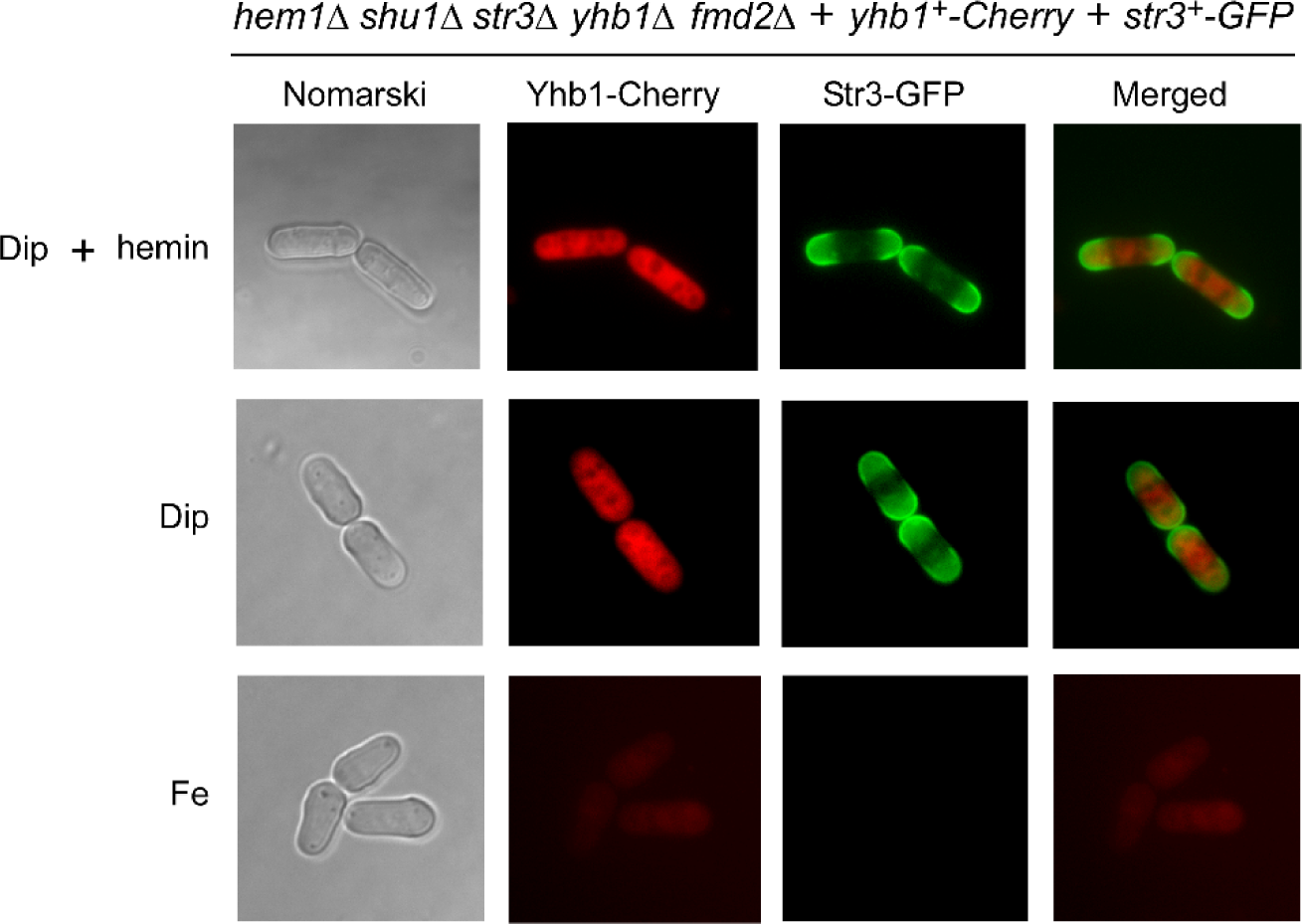
*Localization of Yhb1 and Str3 proteins in iron-starved S. pombe cells deficient in heme biosynthesis. hem1Δ shu1Δ str3Δ yhb1Δ fmd2Δ* cells coexpressing Yhb1-Cherry in combination with Str3-GFP were transferred in ALA-free medium for 6 h. During the final 3 h of culture, aliquots of cells were treated with FeCl_3_ (100 µM), Dip (250 µM) or Dip supplemented with hemin (1 µM). Subsequently, cells were examined by fluorescence microscopy to visualize the localization of Yhb1-Cherry (*center left*) and Str3-GFP (*center right*). The merged images are shown in the far-right panels, whereas Nomarski optics (*far left*) was used to assess cell morphology. Results of microscopy are representative of five independent experiments.

### Yhb1 and Str3 associate with one another to form a protein complex in iron-starved cells

To further investigate whether an association between Yhb1 and Str3 could occur in the presence of hemin under low-iron conditions, mutant cells harboring a *hem1Δ shu1Δ str3Δ yhb1Δ fmd2Δ* quintuple deletion were used to coexpress *yhb1^+^-Cherry* and *str3^+^-GFP* or *yhb1^+^-Cherry* and *GFP* alleles. Each allele was controlled by its respective promoter, except for *GFP*, which was regulated by the thiamine-sensitive *nmt41^+^*promoter. To examine the possibility that Yhb1-Cherry interacts with Str3-GFP, ALA-starved cells coexpressing these proteins were subjected to different treatments: Dip (250 µM) plus hemin (1 µM), Dip alone, and FeCl_3_ (100 µM). After 3 h, whole-cell extracts were prepared and subsequently incubated in the presence of anti-Cherry protein G-Sepharose beads to selectively retain Cherry-tagged Yhb1. Results consistently showed the retention of Yhb1-Cherry on the beads and its presence in the immunoprecipitated fraction (IP) (Fig. 7A). Notably, its detection exhibited a slight enrichment in extracts from cells incubated with hemin under low iron conditions compared to those treated with iron (Fig. 7A). One reason for this is likely due to lower *yhb1^+^* expression in iron-replete conditions (Fig. 1). Importantly, the immunoprecipitated fraction contained Str3-GFP, indicating its interaction with Yhb1-Cherry in extracts from cells grown in the presence of Dip without or with hemin supplementation (Fig. 7A). In contrast, in iron-treated cells where the expression of *str3^+^* gene is repressed (refs), coimmunoprecipitation assays failed to reveal an interaction between Yhb1-Cherry and Str3-GFP (Fig. 7A).

**Fig. 7.**
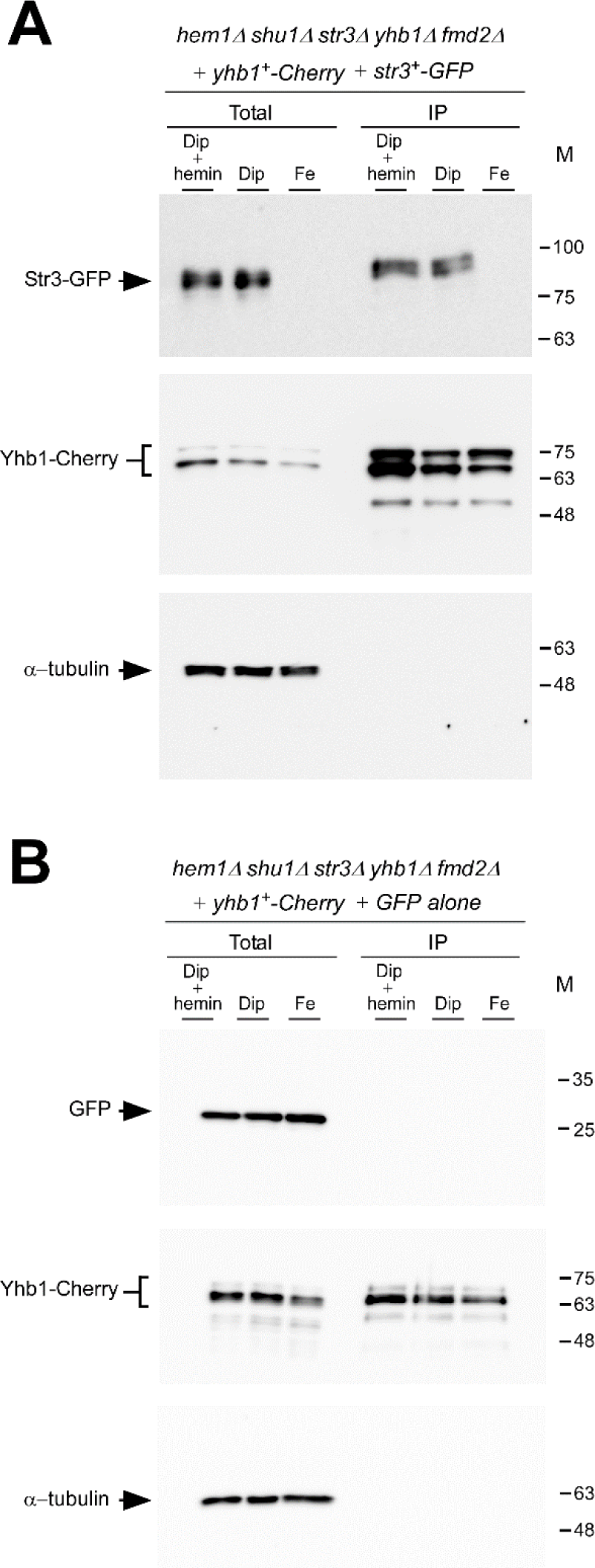
*Yhb1 and Str3 form a heteroprotein complex in iron-deprived cells*. *hem1Δ shu1Δ str3Δ yhb1Δ fmd2Δ* cells coexpressing Yhb1-Cherry in combination with Str3-GFP (*panel A*) or GFP alone (*panel B*) were cultured in an ALA-free medium for 3 h. Subsequently, the ALA-starved cultures were divided into three treatment groups: Dip (250 µM) plus hemin (10 µM), Dip (250 µM), and FeCl_3_ (100 µM) for an additional 3 h prior to cell extract preparations. Whole-cell extracts (Total) were subjected to coimmunoprecipitation (IP) using an anti-Cherry antibody coupled with protein G-Sepharose beads. Bound proteins were eluted from the beads and analyzed by immunoblot assays employing anti-Cherry and anti-GFP antibodies to detect Yhb1-Cherry, Str3-GFP, and GFP alone. Aliquots of total cell extract preparations were included in the assays to confirm the presence of epitope-tagged proteins prior to IP. To serve as an additional control, aliquots of total extracts and bound fractions were probed with an anti-α-tubulin antibody. The molecular weight standards (in kDa) are indicated on the right side of the panels for reference.

To assess the specificity of the coimmunoprecipitation experiments, control cells coexpressing Yhb1-Cherry and GFP alone did not exhibit any interaction between the two proteins in the bound fractions (IP) (Fig. 7B). Further confirmation of the specificity of coimmunoprecipitation assays involved verifying the absence of α-tubulin in the bound fractions (IP) (Fig. 7). Collectively, these findings revealed that Yhb1-Cherry and Str3-GFP associate as a heteroprotein complex when iron-starved cells are exposed to exogenous hemin.

### Mutual interaction of Yhb1-VN and Str3-VC is observed at the cell periphery

To further characterize the interaction between Yhb1 and Str3 in living *S. pombe* cells, we used a bimolecular fluorescence complementation (BiFC) method (Ioannoni et al. 2010; Kerppola 2006; Sung and Huh 2007). Considering the association observed between Yhb1 and Str3 in coimmunoprecipitation assays, we hypothesized that bringing together the nonfluorescent Venus N-terminal (VN) and C-terminal (VC) fragments fused to the C-terminal ends of Yhb1 and Str3, respectively, would reconstitute a functional fluorophore when they are brought together via the Yhb1-Str3 association. Integrative plasmids expressing *yhb1^+^-VN* and *str3^+^-VC* alleles under the control of their own promoters were cotransformed in *hem1Δ shu1Δ str3Δ yhb1Δ fmd2Δ* cells. These cotransformed cells were incubated in an ALA-free medium and subsequently supplemented with FeCl_3_ (100 µM), Dip (250 µM), or a combination of Dip and hemin (1 µM). Examination of the Yhb1-VN-Str3-VC complex’s localization by fluorescence microscopy revealed a BiFC signal at the plasma membrane of cells treated with Dip, and Dip plus hemin (Fig. 8). Although the BiFC signals were not uniformly distributed around the periphery of these cells, specific cell contour regions exhibited heightened fluorescence levels. Conversely, no BiFC signal was detected when Yhb1-VN and Str3-VC were coexpressed in *hem1Δ shu1Δ str3Δ yhb1Δ fmd2Δ* cells treated with iron (Fig. 8), likely due to the repression of *str3^+^* expression under iron-replete conditions (Normant et al. 2018). To verify a protein that produces a fluorescence signal at the plasma membrane, a GFP-tagged *str3^+^* allele was cotransformed with an empty vector in *hem1Δ shu1Δ str3Δ yhb1Δ fmd2Δ* cells. The resulting GFP fluorescence signal localized at the cellular periphery in cells incubated in an ALA-free medium supplemented with Dip, as described previously (Fig. 8) (Normant et al. 2018). As a negative control, *hem1Δ shu1Δ str3Δ yhb1Δ fmd2Δ* cells coexpressing Yhb1-VN and Ctr4-VC proteins. Ctr4 is a cell-surface copper transporter that is unrelated to Yhb1 (Fig. 8) (Ioannoni et al. 2010). As a control, Yhb1-mNeonGreen coexpressed with an empty vector in *hem1Δ yhb1Δ fmd2Δ* cells produced a fluorescence signal in the cytoplasm under Dip, and Dip plus hemin conditions, whereas the signal was weak under iron-replete conditions (Fig. 8). Taken together, these results underscored that the Yhb1-Str3 association is detectable at the plasma membrane of living cells upon exposure to exogenous hemin under low iron conditions.

**Fig. 8.**
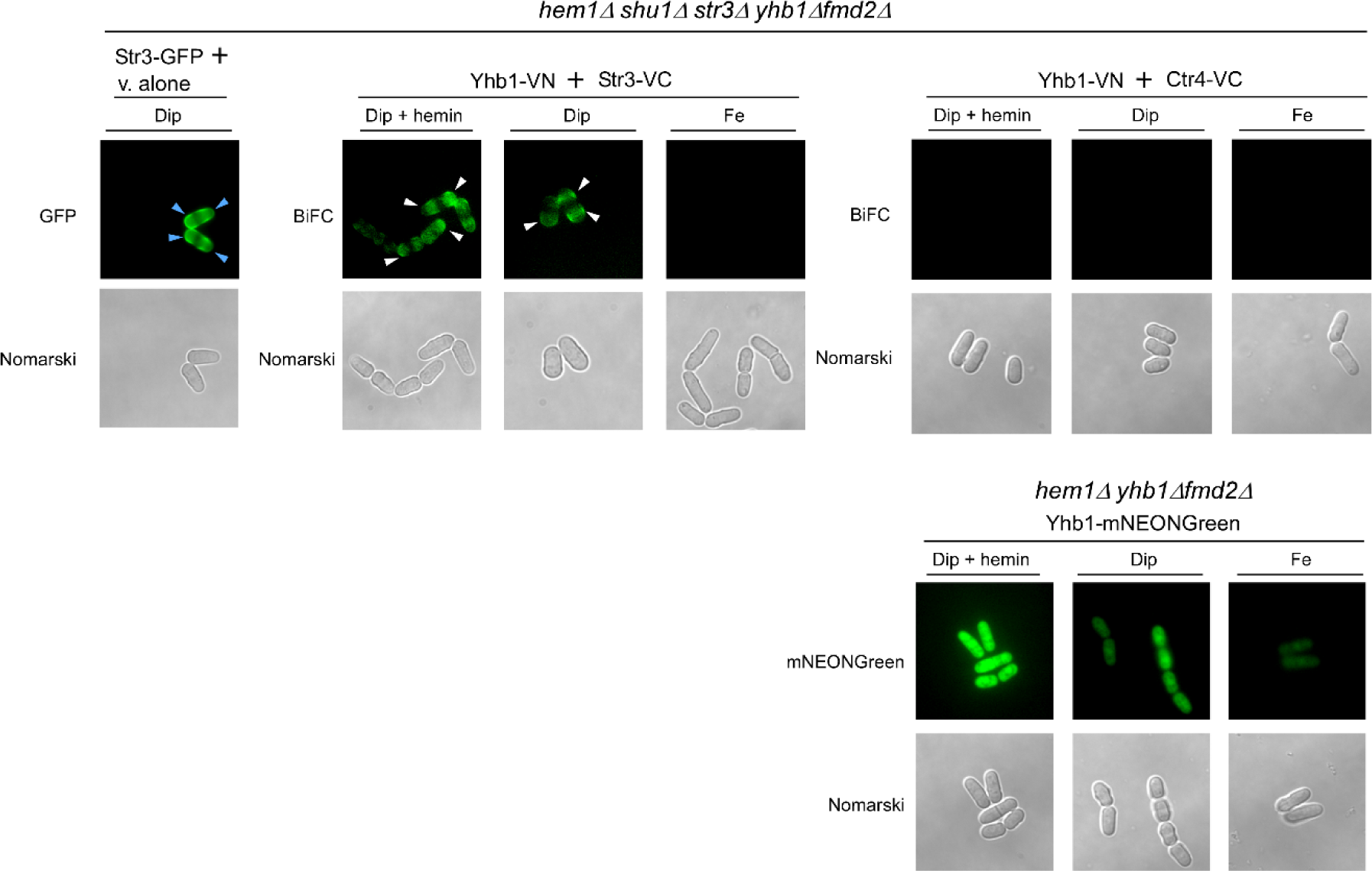
*ALA-starved cells coexpressing Yhb1-VN and Str3-VC generate a BiFC signal at the plasma membrane in the presence of Dip or Dip plus hemin. hem1Δ shu1Δ str3Δ yhb1Δ fmd2Δ* cells expressing Yhb1-VN and Str3-VC proteins were initially incubated in an ALA-free medium for 3 h. Subsequently, these ALA-starved cells were treated with Dip (250 µM) plus hemin (10 µM), Dip (250 µM), or FeCl_3_ (100 µM) for an additional 3 h. Following this incubation period, the cells were examined for BiFC signals using fluorescence microscopy. Exemples of Yhb1VN/Str3VC-mediated BiFC signals at the plasma membrane are denoted by white arrowheads. As a control in parallel experiments, Str3-GFP, which is known to produce a fluorescence signal at the plasma membrane, is indicated by blue arrowheads. The localization of Str3-GFP was assessed using *hem1Δ shu1Δ str3Δ yhb1Δ fmd2Δ* cells coexpressing the *str3^+^-GFP* allele in the presence of an empty vector (v. alone) under the same conditions as mentioned above. As a negative control, *hem1Δ shu1Δ str3Δ yhb1Δ fmd2Δ* cells coexpressing Yhb1-VN and Ctr4-VC proteins. Ctr4 is a cell-surface copper transporter that is unrelated to Yhb1 (*right side*). As a control, Yhb1-mNeonGreen coexpressed with an empty vector in *hem1Δ yhb1Δ fmd2Δ* cells produced a fluorescence signal in the cytoplasm under Dip, and Dip plus hemin conditions, whereas the signal was weak under iron-replete conditions. Cell morphology was monitored using Normarski optics. The results from microscopy represent findings from five independent experiments.

### Str3 is required to protect hem1Δ fmd2Δ mutant cells from nitrosative stress

Based on the results that Yhb1 interacts with the heme transporter Str3, we investigated whether disabling Str3 (*str3Δ*) could increase the sensitivity of *hem1Δ fmd2Δ* cells to DETANONOate when cultured in an ALA-free medium supplemented with hemin under low-iron conditions. First, *hem1Δ fmd2Δ str3Δ* cells were cultured in the presence of ALA to ensure that their growth was comparable to that of *hem1Δ fmd2Δ* and *hem1Δ fmd2Δ yhb1Δ* strains (Fig. 9A). When *hem1Δ fmd2Δ* and *hem1Δ fmd2Δ yhb1Δ* cells (both strains expressing *shu1^+^* and *str3^+^*) were incubated in the absence of ALA and in the presence of 0.25 µM hemin, their cell growth was similar (A_600_ of 6.1 and 6.0 after 47 h) (Fig. 9B). Under these conditions, a *hem1Δ fmd2Δ str3Δ* strain (expressing endogenous *shu1^+^* only) exhibited 73%, 67%, 63%, 48%, and 21% less growth compared to *hem1Δ fmd2Δ* cells after 22, 24, 26, 29, and 47 h, respectively (Fig. 9B). The disruption of the *str3^+^* gene (*str3Δ*) likely reduced the ability of *hem1Δ fmd2Δ str3Δ* cells to grow in the presence of 0.25 µM hemin. We then examined whether exposure to DETANONOate (3 mM) affected the proliferation of the aforementioned strains. In the case of *hem1Δ fmd2Δ* cells (expressing *str3^+^* and *yhb1^+^*), although their ability to grow was diminished (26% growth inhibition after 47 h) compared to untreated *hem1Δ fmd2Δ* cells, they still reached an A_600_ of 4.6 after 47 h (Fig. 9B). When *hem1Δ fmd2Δ yhb1Δ* and *hem1Δ fmd2Δ str3Δ* mutant strains were exposed to DETANONOate, they displayed hypersensitivity to the NO donor compared to these same mutant strains left untreated (Fig. 9B). Remarkably, the poor growth of *hem1Δ fmd2Δ str3Δ* cells (A_600_ of 1.33 after 47 h) was similar to that of *hem1Δ fmd2Δ yhb1Δ* cells (A_600_ of 1.27 after 47 h) (Fig. 9B). Taken together, these results indicated that the loss of function of Str3 phenocopies the effects of Yhb1 disruption by causing hypersensitivity to DETANONOate under hemin-dependent culture conditions.

**Fig. 9.**
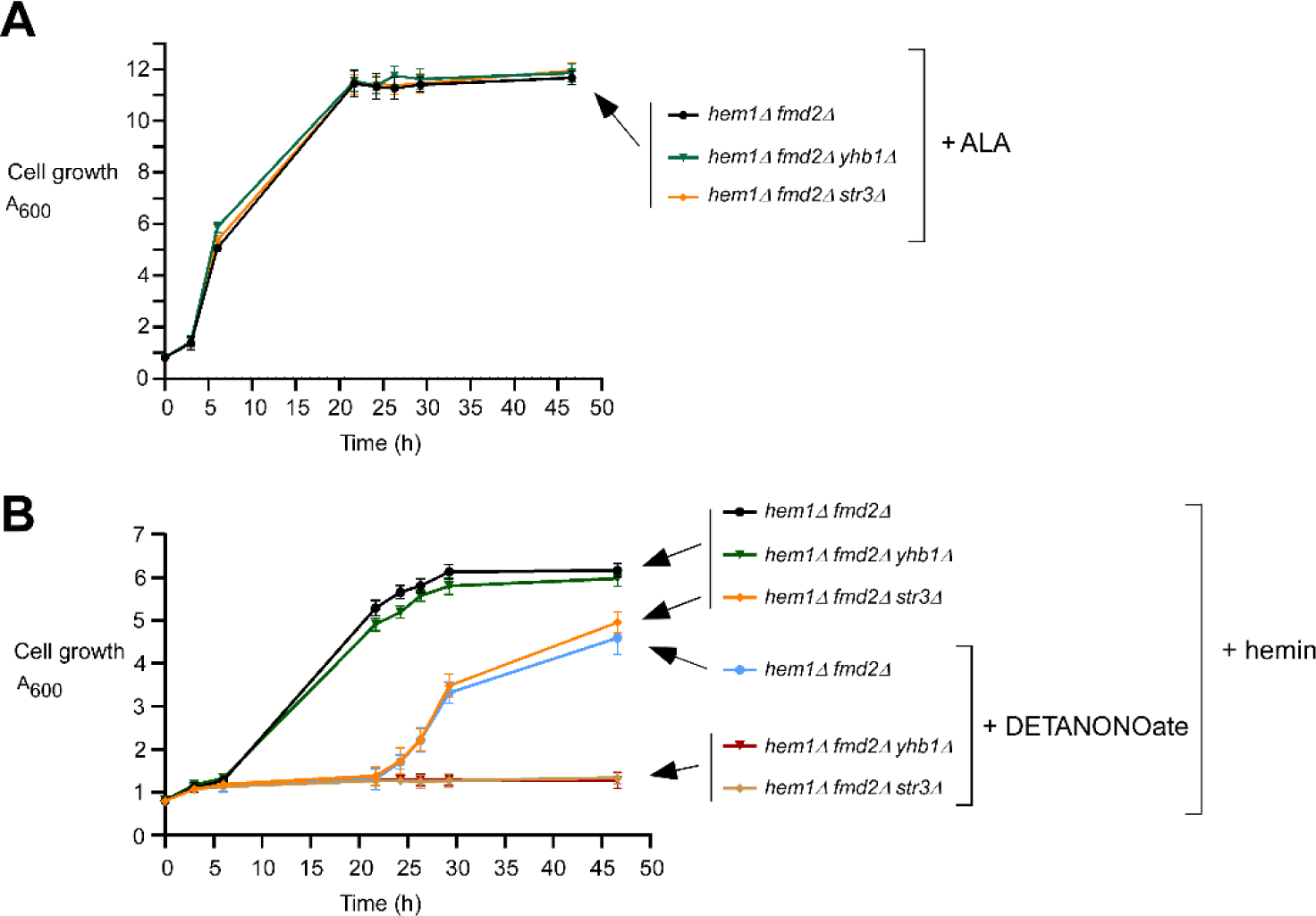
*Loss of the str3^+^ gene mirrors the effect of yhb1^+^ disruption under nitrosative stress conditions*. *A*, Growth of the indicated strains in YES medium that was supplemented with exogenous ALA (200 µM). Strain color codes are as follows: black, *hem1Δ fmd2Δ*; green, *hem1Δ fmd2Δ yhb1Δ*; and, orange, *hem1Δ fmd2Δ str3Δ*. *B*, Cell proliferation assays were performed using the indicated strains grown in ALA-free medium supplemented with hemin (1 µM). Once the cultures reached an A_600_ of 0.8, they were transferred to YES medium (pH 7.2) containing hemin (0.25 µM) without DETANONOate supplementation or with DETANONOate supplementation (3 mM). Growth of the strains was monitored at the specified time points. Strain color codes are as follows: black, *hem1Δ fmd2Δ*; green, *hem1Δ fmd2Δ yhb1Δ*; and, orange, *hem1Δ fmd2Δ str3Δ*. These three strains were left without DETANONOate treatment. Blue, *hem1Δ fmd2Δ*; red, *hem1Δ fmd2Δ yhb1Δ*; and, brown, *hem1Δ fmd2Δ str3Δ*. These last three strains were subjected to DETANONOate treatment (3 mM). The graphs represent the results of three independent experiments performed in biological triplicate, with values presented as averages ± SD.

## Discussion

In our previous study, we reported an interaction between the hemeprotein Yhb1 and the cell-surface heme transporter Str3 using a proximity-dependent biotinylation approach coupled to mass spectrometry (Normant et al. 2021). In our pursuit to further characterize this relationship between these two cellular components, we sought to examine whether they exhibit similarities with respect to their levels of expression in response to changes in iron levels. One rationale for this inquiry stems from the observation that many genes involved in heme acquisition and utilization undergo up-regulation in response to iron deprivation in fungi (Mercier et al. 2008; Rustici et al. 2007; Shakoury-Elizeh et al. 2004). Assessment of the transcript levels of *yhb1^+^* and *str3^+^* confirmed that both genes were upregulated in the presence of the iron chelator Dip compared to their respective levels under basal and iron-replete conditions (Fig. 1). As previously demonstrated for *str3^+^* (Normant et al. 2018), we determined that Fep1 plays a role in the iron-mediated down-regulation of *yhb1^+^* transcription. Notably, the extent of the iron-dependent decrease in *yhb1^+^* mRNA is less pronounced than the effect observed on *str3^+^* mRNA expression. This difference may be due to the positioning of the predicted GATA element in the *yhb1^+^* promoter. The Fep1-binding site is located considerably further upstream, between positions -1443 and -1434 relative to the *yhb1^+^* initiator codon, compared to the *str3^+^* GATA element, which is located between positions - 651 to -642 relative to the *str3^+^* ATG codon. Nevertheless, ChIP analysis revealed that Fep1 is recruited to the *yhb1^+^*promoter under iron-replete conditions in the presence of hemin as the sole source of heme (Fig. 2).

In this study, when heme biosynthesis needed to be blocked, *hem1Δ*-based strains were cultured in the absence of ALA but in the presence of exogenous hemin under low iron conditions. In this case, cells were forced to use their heme uptake machinery. Moreover, in the case of cells treated with DETANONOate, exposure to NO may deactivate the DNA-binding capacity of Fep1, ensuring transcriptional induction of *yhb1^+^* and genes encoding the heme uptake machinery that are subject to Fep1 control. One can envision the following model to interpret a predicted NO-mediated inactivation of Fep1. Studies have shown that the DNA binding activity of Fep1 functions through the binding of a bridging [2Fe-2S]-cluster via its amino acid residues Cys-70, -76, -85 and -88 (Cutone et al. 2016; Hati et al. 2023). Similarly, the *E. coli* transcriptional repressor NsrR requires the presence of an [2Fe-2S]-cluster to bind promoters of its target genes(Tucker et al. 2008; Tucker et al. 2010). Once bound to DNA, NsrR represses transcription by blocking the binding of RNA polymerase. In the presence of NO, a mixture of different dinitrosyl Fe complexes can be formed at the [2Fe-2S]-cluster of NsrR, resulting in a loss of its DNA binding activity, thereby relieving its repression of target genes(Tucker et al. 2008; Tucker et al. 2010). Consistent with the observation that the presence of NO could hinder the binding of Fep1 to chromatin, it has been reported that NO-treated *S. pombe* cells exhibit higher levels of Fe transport gene expression (Astuti et al. 2016). When cells are iron-deficient following treatment with Dip as in our conditions, it is already known that the proteins Grx4 and Fra2 extract the [2Fe-2S]-cluster from Fep1 to inactivate its DNA-binding activity (Hati et al. 2023). By adding DETANONOate treatment to cells already deficient in iron, it would ensure that if the action of Grx4 and Fra2 to strip the [2Fe-2S]-cluster from Fep1 is incomplete, then the presence of NO could ensure the complete inactivation of Fep1 DNA binding activity on promoters of target genes.

It is intriguing that the three fungal NO-responsive transcription factors that have been identified so far are very different from each other, belonging to different families of regulators (Chiranand et al. 2008; Jian et al. 2021; Sarver and DeRisi 2005). In the case of *C. albicans*, the transcription factor Cta4, a Zn_(2)_Cys_(6)_ binuclear cluster regulator, is required for transcriptional induction of *YHB1* gene expression in response to NO (Chiranand et al. 2008). In contrast, the transcription factor Cwt1 acts as a repressor of *C. albicans YHB1* gene under basal conditions (Sellam et al. 2012). In *S. cerevisiae*, the transcription factor that regulates the nitrosative stress response differs from that in *C. albicans*. Induction of *S. cerevisiae YHB1* transcription requires Fzf1, a C_2_H_2_ zinc finger factor that is unrelated to Cta4 (Sarver and DeRisi 2005). In *Fusarium graminearum*, the inducible response to NO requires the presence of the GATA-type transcription factor FgAreB (Jian et al. 2021). This regulatory mechanism relies on the NO-dependent degradation of Fglxr1, a repressor that competes with FgAreB for binding to the SWI/SNF complex. Nitrosative stress triggers the degradation of Fglxr1 by the proteasome machinery, facilitating the recruitment of the SWI/SNF complex by FgAreB, ultimately leading to the activation of gene expression in response to NO (Jian et al. 2021). At this time, it is unclear how *S. pombe* detects the presence of NO and which specific signaling pathways mediate responses to NO and reactive nitrogen intermediates (RNI). While transcription factor candidates have been proposed to act as NO-responsive master regulators to promote a comprehensive nitrosative stress defense mechanism, none of them, upon their inactivation, fully prevents the induction of gene transcription in response to nitrosative stress. Therefore, the identity of the *S. pombe* NO-responsive major transcription factor remains unresolved.

When heme biosynthesis is selectively blocked (*hem1Δ*) and ALA is absent under iron starvation conditions, *S. pombe* cells must utilize at least one of the two heme uptake systems to survive in the presence of exogenous hemin. In this study, we used cells lacking the heme receptor Shu1, thereby restricting them to rely on Str3 for heme assimilation. Under these conditions, our previous studies have demonstrated that Str3 can interact with the peroxiredoxin Tpx1 and the sulfiredoxin Srx1 (Normant et al. 2021; Vahsen et al. 2023). Similarly, in this study, co-immunoprecipitation and BiFC experiments revealed the interaction between Yhb1-Cherry and Str3-GFP proteins in iron-starved cells grown in the presence of hemin. These results raise the question of whether one or the other interacting partner of Str3 could be favored to interact with Str3 under nitrosative stress conditions. When cells are exposed to NO, there is an emergence of reactive nitrogen intermediates and reactive oxygen species (ROS) as secondary metabolites within the cells. To counteract this emergence of ROS, a substantial portion of the Tpx1 pool is predicted to undergo a transition from a heme-binding protein to an antioxidant protein. When Tpx1 functions as a scavenger of hydrogen peroxide or alkyl hydroperoxides, it loses its propensity to bind to heme (Watanabe et al. 2017). In the case of Srx1, its primary role is to reactivate hyperoxidized Tpx1 by reducing it to its sulfinic form, thereby restoring Tpx1’s competence in oxidative stress defense(Rhee et al. 2007; Rhee and Woo 2011). The sulfinic reductase activity of Srx1 that is required for reactivating hyperoxidized Tpx1 is ATP-dependent. When Srx1 binds heme, it is predicted to inhibit its ability to bind ATP, thereby abolishing its sulfinic reductase activity(Vahsen et al. 2023). When ROS production increases, a majority of the Srx1 pool is predicted to primarily serve in the peroxidase catalytic cycle, aiming to reactivate hyperoxidized Tpx1. During nitrosative stress, which induces the emergence of ROS, Tpx1 and Srx1 fulfill their antioxidant roles, rendering them unavailable to form complexes with heme. Consequently, it is expected that there will be a significant preference for the Yhb1-Str3 interaction, enabling Yhb1 to mobilize heme from Str3 in the presence of NO.

Other cellular heme-binding proteins play a role in maintaining a protected intracellular pool of heme available for distribution to appropriate client proteins and compartments (Swenson et al. 2020). Among them, glyceraldehyde-3-phosphate dehydrogenase (GAPDH) is known for mobilizing cytosolic heme and delivering it to client proteins, including soluble guanylyl cyclase β (sGCβ) and the heme-dependent nuclear transcription factor Hap1 (Dai et al. 2020; Sweeny et al. 2018). However, studies have shown that exposure to nitrosative stress impairs GAPDH’s heme-binding ability (Chakravarti et al. 2010; Hanna et al. 2016). This unfavorable condition for GAPDH would lead to a strong bias toward heme delivery to Yhb1, as GAPDH becomes unavailable to form complexes with heme.

In this study, we examined the impact of three mutant variants of Yhb1, denoted Yhb1^M1^, Yhb1^M2^, and Yhb1^M3^. These mutants were designed to substitute amino acid residues that are highly conserved among flavoHbs and may serve as potential heme ligands(El Hammi et al. 2012). The Yhb1^M1^ mutant harbored a single mutation (His^114^A) at the predicted heme coordination site. Results showed that cells expressing Yhb1^M1^-Cherry exhibited hemin-dependent growth deficiency in the presence of DETANONOate, but supported hemin-dependent growth in its absence. We concluded that the Yhb1 His^114^ residue is essential for the NO dioxygenase (NOD) activity of Yhb1 under nitrosative stress conditions in heme synthesis-deficient cells grown in the presence of hemin. Similar outcomes were observed with the Yhb1^M2^ and Yhb1^M3^ mutants. It is important to note that we did not directly assess the NOD activity of the mutant derivatives of Yhb1. Instead, we inferred potential losses in NOD function based on existing knowledge concerning the impact of different highly conserved amino acid residues on flavoHb activity in homologous enzymes present in bacterial and fungal species(El Hammi et al. 2012; Ilari and Boffi 2008).

Non-iron metalloporphyrins (MPPs) are taken up by bacteria and yeasts via heme uptake systems, and they have antibacterial/antifungal activity that is thought to result from interference with essential hemeproteins(Hu et al. 2013; Roy et al. 2022; Stojiljkovic et al. 2001). In the present study, we used the non-iron metalloporphyrin ZnMP instead of hemin. Moreover, we used a medium supplemented with a minimal concentration of ALA capable of sustaining minimal yeast growth of *hem1Δ*-based strains. Under these conditions, we ensured that ZnMP was present in excess (10-fold) to be efficiently incorporated into Yhb1 rather than heme. Results showed that when ZnMP was acquired via the Str3-dependent pathway, it impeded Yhb1 function, rendering cells hypersensitive to DETANONOate. Hence, from a practical application standpoint, the use of toxic MPPs or similar compounds recognized by a fungal uptake system may represent a promising strategy for combating fungal resistance to nitrosative stress displayed by both saprophytic and pathogenic yeasts.

## Experimental procedures

### Strains and growth conditions

This study utilized *S. pombe* strains detailed in Table 1. The strains were cultured in yeast extract with supplements (YES) medium under non-selective growth conditions, following previously established protocols (Sabatinos and Forsburg 2010). In the case of strains lacking the *hem1^+^* gene (*hem1Δ*), they were maintained alive with ALA supplementation (200 µM). Alternatively, to ensure viability, YES medium for *hem1Δ* strains was supplemented with exogenous hemin (1 µM), forcing the strains to uptake exogenous hemin for growth. Strains requiring plasmid integration or transformation were grown on synthetic Edinburgh minimal medium (EMM) lacking specific amino acids to select the integrated DNA fragment or plasmid. In experiments involving YES medium with the NO donor DETANONOate (Cayman Chemicals, Ann Arbor, MI), the pH was adjusted to 7.2 using HEPES buffer (1 M HEPES-NaOH, pH 7.5). DETANONOate solution, freshly prepared on ice in the presence of 0.01 M NaOH, was added to buffered YES at a final concentration of 3 mM, unless specified otherwise.

**Table 1.** – S. pombe strains used in this study

| Strain | Genotype | Source or reference |
| --- | --- | --- |
| FY435 | *h<sup>+</sup> his7-366 leu1-32 ura4-Δ18 ade6-M210* | Mourer *et al.* 2015 |
| TMY1 | *h<sup>+</sup> his7-366 leu1-32 ura4-Δ18 ade6-M210 hem1Δ::KAN<sup>r</sup>* | Mourer *et al.* 2015 |
| TVY1 | *h<sup>+</sup> his7-366 leu1-32 ura4-Δ18 ade6-M210 hem1Δ::loxP fep1Δ::KAN<sup>r</sup>* | Vahsen *et al.* 2023 |
| FLYPY1 | *h<sup>+</sup> his7-366 leu1-32 ura4-Δ18 ade6-M210 hem1Δ::loxP fmd2Δ::KAN<sup>r</sup>* | This study |
| FLYPY2 | *h<sup>+</sup> his7-366 leu1-32 ura4-Δ18 ade6-M210 hem1Δ::loxP yhb1Δ::loxP* | This study |
| FLYPY3 | *h<sup>+</sup> his7-366 leu1-32 ura4-Δ18 ade6-M210 hem1Δ::loxP yhb1Δ::loxP fmd2Δ::KAN<sup>r</sup>* | This study |
| FLYPY4 | *h<sup>+</sup> his7-366 leu1-32 ura4-Δ18 ade6-M210 hem1Δ::loxP shu1Δ::loxP fmd2Δ::KAN<sup>r</sup>* | This study |
| FLYPY5 | *h<sup>+</sup> his7-366 leu1-32 ura4-Δ18 ade6-M210 hem1Δ::loxP str3Δ::loxP fmd2Δ::KAN<sup>r</sup>* | This study |
| FLYPY6 | *h<sup>+</sup> his7-366 leu1-32 ura4-Δ18 ade6-M210 hem1Δ::loxP shu1Δ::loxP str3Δ::loxP yhb1Δ::loxP fmd2Δ::loxP* | This study |
| FLYPY7 | *h<sup>+</sup> his7-366 leu1-32 ura4-Δ18 ade6-M210 hem1Δ::loxP shu1Δ::loxP str3Δ::loxP yhb1Δ::loxP* | This study |

For yeast liquid proliferation assays, precultures of specified strains were initially grown in YES medium containing ALA (200 µM) and Dip (250 µM). At this stage, the presence of ALA ensured heme biosynthesis, whereas the iron chelator Dip induced transcription of iron-starvation responsive genes like *yhb1^+^* and *str3^+^*. Upon reaching mid-logarithmic phase, cells were washed twice, diluted 1000-fold in YES containing Dip (25 µM or the specified concentration), and supplemented with hemin (1 µM) without ALA, unless stated otherwise. Cultures were then initiated until reaching an A_600_ of 0.8. At this cell density, cultures were transferred in YES buffered to pH 7.2 and then incubated without DETANONOate or with DETANONOate treatment. At this stage (zero time point), cell growth was monitored by measuring optical density at A_600_.

### Plasmids

The plasmid pJK-1679yhb1prom was constructed as follows: a SacII-BamHI PCR-amplified fragment from the *yhb1^+^*promoter, encompassing the initial 1,679 bp of the 5’-untranslated region, was inserted into the SacII and BamHI sites of pJK148 (Keeney and Boeke 1994). For the creation of the integrative pJK-1679yhb1^+^ plasmid, a 1,284-bp BamHI-SalI-amplified DNA fragment containing the *yhb1+* gene was cloned into the corresponding sites of pJK-1679yhb1prom. The same *yhb1^+^* DNA fragment was amplified without its stop codon and then cloned into the BamHI and SalI sites of pJK-1679yhb1prom, resulting in the formation of the plasmid pJK-1679yhb1nostop.

Subsquently, the TAP, Cherry, mNeonGreen, and VN coding sequences were isolated by PCR from pJK-1478NTAPfep1^+^ (Pelletier et al. 2005), pJK148mfc1^+^-Cherry (Beaudoin et al. 2011), pFA6a-NeonGreen-KanMX6 (Addgene plasmid 129099), and pSP1ctr5^+^-VN (Ioannoni et al. 2010), respectively. Each PCR product was generated using primer sets that introduced SalI and KpnI sites at the ends of the PCR products. After purification and digestion with SalI and KpnI, these PCR products were inserted in-frame with *yhb1^+^* into the corresponding sites of pJK-1679yhb1nostop, resulting in the generation of four plasmids, denoted as pJKyhb1^+^-TAP, pJKyhb1^+^-Cherry, pJKyhb1^+^-mNeonGreen, and pJKyhb1^+^-VN, respectively. The construction of the integrative plasmids pBPade-785str3^+^-GFP, pBPnmt41x-GFP, and pBPade-785str3^+^-VC has been previously described (Brault et al. 2022; Normant et al. 2021; Normant et al. 2018).

The plasmid pJKyhb1^+^-Cherry was used to introduce mutations in the coding sequence of *yhb1^+^*. Specifically, in the *yhb1^M1^-Cherry* mutant allele, the codon corresponding to His^114^ was replaced with a nucleotide triplet encoding an alanine residue. Similarly, the *yhb1^M2^-Cherry* mutant allele exhibited substitutions where the codons for Tyr^58^, His^114^, Tyr^124^, Pro^125^, Ile^126^, and Val^127^ were replaced by nucleotide triplets encoding alanine residues. In the case of the *yhb1^M3^-Cherry* mutant allele, it harbored the same substitutions as *yhb1^M2^-Cherry*, with an additional substitution where the codon corresponding to Glu^167^ was replaced by a DNA triplet encoding alanine. These site-specific mutations were created using a PCR overlap extension method (Ho et al. 1989).

### RNA extraction and mRNA expression analysis

Total RNA was isolated from cell cultures using the hot phenol method, as previously described (Chen et al. 2003). For the analysis of *yhb1^+^*mRNA, two different methods were used to assess its expression. Initially, *yhb1^+^*transcripts were analyzed using RNase protection assays as previously described (Mercier et al. 2008). To monitor *yhb1^+^* mRNA levels, the pSK*yhb1^+^* plasmid was designed to generate a riboprobe specifically hybridizing to *yhb1^+^* transcripts. Plasmid pSK*yhb1^+^* was created by inserting a 187-bp BamHI-EcoRI fragment from the *yhb1^+^* gene into the corresponding restriction sites of pBluescript SK. This plasmid was designed to produce an antisense RNA probe that specifically hybridizes to the region between positions +108 to +295 of *yhb1^+^*. In the case of the antisense *act1^+^* riboprobe, it was produced from plasmid pSK*act1^+^* (Peter et al. 2008), following the protocol previously described (Mercier et al 2008). The detection of *act1^+^*transcript served as an internal control for quantification of the RNase protection products.

Subsequently, *yhb1^+^*, *str3^+^*, *act1^+^*, and *tub1^+^* transcripts were subjected to real-time quantitative reverse transcription PCR (RT-qPCR) assays, as previously described (Brault et al. 2022). Reverse transcription reactions, cDNA synthesis, and qPCR reactions followed the procedures outlined as previously described (Brault et al. 2022). Each target transcript was assessed in experiments involving a minimum of three biological replicates. Fold changes for each transcript were determined using the ΔΔCt method, normalized to *act1^+^* or *tub1^+^*, which served as internal controls (Livak and Schmittgen 2001; Protacio et al. 2022; Schmittgen and Livak 2008). The calculations were performed using the following equation: ΔΔCt = [(Ct gene–Ct ref) in iron-starved cells] versus [(Ct gene–Ct ref) in untreated or iron-replete cells], where ref was *act1^+^*, or ΔΔCt = [(Ct gene–Ct ref) in untreated cells] versus [(Ct gene–Ct ref) in DETANONOate-treated cells], where ref was *tub1^+^*. To monitor *yhb1^+^* transcript levels by RT-qPCR, a primer pair was utilized, allowing the detection of an amplicon corresponding to the coding region between positions +290 and +398, down to the first nucleotide of the *yhb1^+^* initiator codon. For the detection of *str3^+^* expression, a primer pair was used for amplifying the coding sequence between +585 and +690, down to the first base of the ATG codon of *str3^+^*. Regarding *act1^+^* and *tub1^+^*, the amplicons corresponded to the coding regions between positions +173 to +280 and +516 to +626, respectively.

### ChIP assays

To further examine whether Fep1 directly interacts with the *yhb1^+^*promoter, *hem1Δ php4Δ fep1Δ* cells expressing untagged or TAP-tagged Fep1 were grown until reaching an A_600_ of 0.35. Subsequently, they were transferred to an ALA-free YES medium supplemented with FeCl_3_ (75 µM) for 5 h. After washings, the cells were transferred to ALA- free medium supplemented with hemin (5 µM) and treated with Dip (250 µM) or FeCl_3_ (100 µM). Following a 3-h treatment, *in vivo* cross-linking of proteins was performed by incubating cell cultures with 1% formaldehyde for 20 min. After formaldehyde-mediated cross-links and neutralization with glycine, cell lysates were prepared by glass bead disruption, as previously described (Brault et al. 2016; Larochelle et al. 2012). Samples were subsequently sonicated to shear chromatin DNA into fragments approximately ranging from 500 to 1000 bp. Immunoprecipitation of TAP-Fep1 bound to chromatin was performed using immunoglobin G (IgG)-Sepharose beads, following the procedures as previously described (Jacques et al. 2014). Bead manipulation, which includes washings, elution, reversal of cross-linking, and DNA precipitation, was performed in accordance with the protocols previously described (Adam et al. 2001; Jbel et al. 2009). Quantification of immunoprecipitated DNA was conducted by quantitative real-time PCR (qPCR) using different sets of primers spanning *yhb1^+^* and *shu1^+^* promoter regions. TAP-tagged Fep1 occupancy at the *yhb1^+^* or *shu1^+^* locus was calculated as the enrichment of specific genomic *yhb1^+^* and *shu1^+^* promoter regions relative to a GATA-free 18S ribosomal DNA coding region used as an internal background control. The primers used for amplifying the *yhb1^+^* promoter region were denoted yhb1-1486 (5’-TCAACCTTACATTGACTAGGTTCTT-3’) and yhb1-1385 (5’-GGACTGTGGTTAAAGTGGATCA-3’). The two sets of primers used for amplifying the *shu1^+^* promoter region and a 18S ribosomal DNA coding region were previously described (Brault et al. 2016; Mourer et al. 2015). Each qPCR run was performed in triplicate using the Perfecta SYBR Green Fast mix (Quanta) on a CFX96 Touch Real-Time PCR instrument (BioRad). All ChIP experiments were repeated at least three times using independent chromatin preparations.

### Protein extraction, Western blot and coimmunoprecipitation assays

To analyze the steady-state levels of Yhb1 protein upon exposure to DETANONOate under low iron conditions, whole cell extracts were prepared using glass beads and a FastPrep-24 instrument (MP Biomedicals, Solon, OH). Cells were lysed in TMN_150_ buffer containing 50 mM Tris-HCl (pH 7.5), 150 mM NaCl, 5 mM MgCl_2_, 1% Nonidet P-40, 1 mM phenylmethylsulfonyl fluoride (PMSF), and a complete protease inhibitor cocktail (P8340, Sigma-Aldrich). Equal amounts of cell lysates were resolved on 8% sodium dodecyl sulfate (SDS)-polyacrylamide gels, and the proteins were then transferred to nitrocellulose membranes via electroblotting. For quantitative Western blot assays, Yhb1-TAP and α-tubulin proteins were detected using the polyclonal anti-mouse IgG antibody (ICN Biomedicals) and monoclonal anti-α-tubulin antibody (clone B-5-1-2; Sigma-Aldrich), respectively. Following incubation with primary antibodies, the membranes were washed and incubated with IRDye® 800CW conjugated donkey anti-rabbit IgG secondary antibody and DyLight 680-conjugated goat anti-mouse secondary antibody. Subsequently, the membranes were probed using an Odyssey infrared imaging system (LI-COR) for quantification purposes.

To investigate the interaction between Yhb1 and Str3 in *S. pombe*, *hem1Δ shu1Δ str3Δ yhb1Δ fmd2Δ* cells were cotransformed with integrative plasmids pJKyhb1^+^-Cherry and pBPade-785str3^+^-GFP or pJKyhb1^+^-Cherry and pBPnmt41x-GFP. Cotransformed cells were grown until reaching an A_600_ of 0.5. At this point, cells were transferred to an ALA-free medium for 5 h and subjected to three experimental conditions: presence of hemin (5 µM) and Dip (250 µM); presence of Dip only; and, presence of FeCl_3_ (100 µM). After 3 h, total cell lysates were prepared by glass bead disruption using a FastPrep-24 instrument. Coimmunoprecipitation assays were performed as previously described (Mourer et al. 2019), except that total extracts were subjected to pull-down using monoclonal antibodies against Cherry bound to protein G-Sepharose 4 Fast Flow beads (GE Healthcare). The suspensions were end-over-end mixed on a rotating wheel for 2 h at 4°C. The beads were washed four times with TMN_150_ buffer and then transferred to fresh microtubes prior to a final wash. The immunoprecipitates were resuspended in an SDS loading buffer (240 mM Tris-HCl, pH 6.8, 40% glycerol, 8% SDS, 2.8 mM β-mercaptoethanol, 1 mM DTT, 6 M urea, and 2 M thiourea) and proteins were dissociated from the beads by heating at 65°C for 30 min. The resulting immunoprecipitated proteins were resolved by electrophoresis on 10% SDS-polyacrylamide gels and analyzed by immunoblot assays. Immunodetection of Cherry-tagged Yhb1, GFP-tagged Str3, and α-tubulin was carried out using the following primary antibodies: polyclonal anti-mCherry antibody E5D8F (Cell Signaling Technology), monoclonal anti-GFP antibody B-2 (Santa Cruz Biotechnology), and monoclonal anti-α-tubulin antibody (clone B-5-1-2, Sigma-Aldrich). After incubation with primary antibodies, the Western blot membranes were washed and incubated with appropriate horseradish peroxidase-conjugated secondary antibodies (Amersham Biosciences). The membranes were developed using enhanced chemiluminescence reagents and visualized by chemiluminescence using an ImageQuant LAS 4000 instrument (GE Healthcare) equipped with a Fujifilm High Sensitivity F0.85 43 mm camera.

### Fluorescence microscopy and BiFC analysis

Fluorescence and differential interference contrast images (Nomarski) of cells were acquired using a Nikon Eclipse E800 epifluorescence microscope (Nikon, Melville, NY) outfitted with a Hamamatsu ORCA-ER digital cooled camera (Hamamatsu, Bridgewater, NJ). The cells were viewed using 1000× magnification with two filters: 465-495 nm (for GFP and mNeonGreen signals) and 510-560 nm (for Cherry signal). Representative cell fields depicted in this study are derived from a minimum of five independent experiments. Furthermore, the displayed cell fields represent protein localization in 200 cells examined per condition.

For fluorescence microscopic visualization of ZnMP accumulation, liquid cultures of the specified strains were grown until reaching an A_600_ of 0.8. At this cell density, cultures were transferred in YES buffered to pH 7.2 and then incubated in a medium supplemented with a minimal concentration of ALA (1 µM) to sustain minimal yeast growth, along with a combination of ZnMP (10 µM) either without DETANONOate supplementation or with DETANONOate supplementation (3 mM) for 16 h. Under these growth conditions, we ensured that ZnMP was present in excess to be efficiently incorporated into hemeproteins. ZnMP accumulation was stopped by adding 5 volumes of ice-cold 5% BSA in PBS. After centrifugation, cells were resuspended in ice-cold 2% BSA in PBS and examined by fluorescence microscopy.

### Data availability

All data are included in the present manuscript. Strains and plasmids used for this study are available upon request. The authors state that all results obtained for confirming the conclusions presented in the article are represented fully within the article.

## Acknowledgments

T.V. is recipient of a studentship from the Fonds de Recherche du Québec (FRQ-S). This study was supported by the Canadian Institutes of Health Research (CIHR, grant #202303PJ4-501788-MID-CFDA-36776) to S.L.

## Conflict of interest

The authors declare that they have no conflict of interest with the content of this article.

## Author contributions

Conceptualization, F.L.Y.P., T.V., A.B., R.N., and S.L.; Methodology, F.L.Y.P., T.V., A.B., R.N., and S.L.; Validation, F.L.Y.P., T.V., A.B., R.N., and S.L.; Formal Analysis, F.L.Y.P., T.V., A.B., R.N., and S.L.; Investigation, F.L.Y.P., T.V., A.B., R.N., and S.L.; Resources, F.L.Y.P., T.V., A.B., and S.L.; Writing – original draft preparation F.L.Y.P. and S.L.; Review and editing, F.L.Y.P., T.V., A.B., R.N., and S.L.; Supervision, T.V., A.B., and S.L.; Project administration, S.L.; Funding acquisition, S.L. All authors have read, reviewed and approved the final version of the manuscript.

## Notes

### Competing Interest Statement

The authors have declared no competing interest.

## References.

Adam M, Robert F, Larochelle M, Gaudreau L (2001) H2A.Z is required for global chromatin integrity and for recruitment of RNA polymerase II under specific conditions. Mol. Cell. Biol. 21:6270–6279 doi: 10.1128/MCB.21.18.6270-6279.2001.

Astuti RI, Watanabe D, Takagi H (2016) Nitric oxide signaling and its role in oxidative stress response in *Schizosaccharomyces pombe*. Nitric oxide 52:29–40 doi:10.1016/j.niox.2015.11.001

Beaudoin J, Ioannoni R, Lopez-Maury L, et al. (2011) Mfc1 is a novel forespore membrane copper transporter in meiotic and sporulating cells. J. Biol. Chem. 286:34356–34372 doi:10.1074/jbc.M111.280396

Bonamore A, Boffi A (2008) Flavohemoglobin: structure and reactivity. IUBMB life 60:19–28 doi:10.1002/iub.9

Brault A, Mbuya B, Labbé S (2022) Sib1, Sib2, and Sib3 proteins are required for ferrichrome-mediated cross-feeding interaction between *Schizosaccharomyces pombe* and *Saccharomyces cerevisiae*. Front. Microbiol. 13:962853 doi:10.3389/fmicb.2022.962853

Brault A, Rallis C, Normant V, Garant JM, Bahler J, Labbé S (2016) Php4 is a key player for iron economy in meiotic and sporulating cells. G3 6:3077–3095 doi:10.1534/g3.116.031898

Chakravarti R, Aulak KS, Fox PL, Stuehr DJ (2010) GAPDH regulates cellular heme insertion into inducible nitric oxide synthase. Proc. Natl. Acad. Sci. USA 107:18004–9 doi:10.1073/pnas.1008133107

Chambers IG, Willoughby MM, Hamza I, Reddi AR (2021) One ring to bring them all and in the darkness bind them: The trafficking of heme without deliverers. Biochim. Biophys. Acta Mol. Cell Res. 1868:118881 doi:10.1016/j.bbamcr.2020.118881

Chen D, Toone WM, Mata J, et al. (2003) Global transcriptional responses of fission yeast to environmental stress. Mol. Biol. Cell 14:214–229 doi:10.1091/mbc.E02-08-0499

Chiranand W, McLeod I, Zhou H, et al. (2008) CTA4 transcription factor mediates induction of nitrosative stress response in *Candida albicans*. Eukaryot. Cell 7:268–78 doi:10.1128/ec.00240-07

Cutone A, Howes BD, Miele AE, et al. (2016) *Pichia pastoris* Fep1 is a [2Fe-2S] protein with a Zn finger that displays an unusual oxygen-dependent role in cluster binding. Sci. Rep. 6:31872 doi:10.1038/srep31872

Dai Y, Sweeny EA, Schlanger S, Ghosh A, Stuehr DJ (2020) GAPDH delivers heme to soluble guanylyl cyclase. J. Biol. Chem. 295:8145–8154 doi:10.1074/jbc.RA120.013802

de Jesús-Berríos M, Liu L, Nussbaum JC, Cox GM, Stamler JS, Heitman J (2003) Enzymes that counteract nitrosative stress promote fungal virulence. Curr. Biol. 13:1963–8 doi:10.1016/j.cub.2003.10.029

Donegan RK, Moore CM, Hanna DA, Reddi AR (2019) Handling heme: The mechanisms underlying the movement of heme within and between cells. Free Radic. Biol. Med. 133:88–100 doi:10.1016/j.freeradbiomed.2018.08.005

Dutt S, Hamza I, Bartnikas TB (2022) Molecular mechanisms of iron and heme metabolism. Ann. Rev. Nutr. 42:311–335 doi:10.1146/annurev-nutr-062320-112625

El Hammi E, Warkentin E, Demmer U, Marzouki NM, Ermler U, Baciou L (2012) Active site analysis of yeast flavohemoglobin based on its structure with a small ligand or econazole. FEBS J. 279:4565–75 doi:10.1111/febs.12043

Glorieux C, Calderon PB (2017) Catalase, a remarkable enzyme: targeting the oldest antioxidant enzyme to find a new cancer treatment approach. Biol. Chem. 398:1095–1108 doi:10.1515/hsz-2017-0131

Hanna DA, Harvey RM, Martinez-Guzman O, et al. (2016) Heme dynamics and trafficking factors revealed by genetically encoded fluorescent heme sensors. Proc. Natl. Acad. Sci. USA 113:7539–44 doi:10.1073/pnas.1523802113

Hati D, Brault A, Gupta M, et al. (2023) Iron homeostasis proteins Grx4 and Fra2 control activity of the Schizosaccharomyces pombe iron repressor Fep1 by facilitating [2Fe-2S] cluster removal. J. Biol. Chem. 299:105419 doi:10.1016/j.jbc.2023.105419

Ho SN, Hunt HD, Horton RM, Pullen JK, Pease LR (1989) Site-directed mutagenesis by overlap extension using the polymerase chain reaction. Gene 77:51–59

Honsa ES, Maresso AW, Highlander SK (2014) Molecular and evolutionary analysis of NEAr-iron Transporter (NEAT) domains. PloS One 9:e104794 doi:10.1371/journal.pone.0104794

Hu G, Caza M, Cadieux B, Chan V, Liu V, Kronstad J (2013) Cryptococcus neoformans requires the ESCRT protein Vps23 for iron acquisition from heme, for capsule formation, and for virulence. Infect. Immun. 81:292–302 doi:10.1128/IAI.01037-12; 10.1128/IAI.01037-12

Ilari A, Boffi A (2008) Structural studies on flavohemoglobins. Methods Enzymol. 436:187–202 doi:10.1016/s0076-6879(08)36010-8

Ioannoni R, Beaudoin J, Mercier A, Labbé S (2010) Copper-dependent trafficking of the Ctr4-Ctr5 copper transporting complex. PloS One 5:e11964 doi:10.1371/journal.pone.0011964

Jacques JF, Mercier A, Brault A, Mourer T, Labbé S (2014) Fra2 is a co-regulator of Fep1 inhibition in response to iron starvation. PloS One 9:e98959 doi:10.1371/journal.pone.0098959; 10.1371/journal.pone.0098959

Jbel M, Mercier A, Pelletier B, Beaudoin J, Labbé S (2009) Iron activates in vivo DNA binding of *Schizosaccharomyces pombe* transcription factor Fep1 through its amino-terminal region. Eukaryot. Cell 8:649–664 doi:10.1128/EC.00001-09

Jian Y, Liu Z, Wang H, et al. (2021) Interplay of two transcription factors for recruitment of the chromatin remodeling complex modulates fungal nitrosative stress response. Nature Comm. 12:2576 doi:10.1038/s41467-021-22831-8

Keeney JB, Boeke JD (1994) Efficient targeted integration at leu1-32 and ura4-294 in *Schizosaccharomyces pombe*. Genetics 136:849–856

Kerppola TK (2006) Visualization of molecular interactions by fluorescence complementation. *Nature Rev*. Mol. Cell Biol. 7:449–456 doi:10.1038/nrm1929

Kumar S, Bandyopadhyay U (2005) Free heme toxicity and its detoxification systems in human. Toxicol. Lett. 157:175–88 doi:10.1016/j.toxlet.2005.03.004

Labbé S, Mourer T, Brault A, Vahsen T (2020) Machinery for fungal heme acquisition. Curr. Genet. 66:703–711 doi:10.1007/s00294-020-01067-x

Larochelle M, Lemay JF, Bachand F (2012) The THO complex cooperates with the nuclear RNA surveillance machinery to control small nucleolar RNA expression. Nucleic Acids Res. 40:10240–10253 doi:10.1093/nar/gks838; 10.1093/nar/gks838

Liu L, Hausladen A, Zeng M, Que L, Heitman J, Stamler JS (2001) A metabolic enzyme for S-nitrosothiol conserved from bacteria to humans. Nature 410:490–4 doi:10.1038/35068596

Liu L, Zeng M, Hausladen A, Heitman J, Stamler JS (2000) Protection from nitrosative stress by yeast flavohemoglobin. Proc. Natl. Acad. Sci. USA 97:4672–6 doi:10.1073/pnas.090083597

Livak KJ, Schmittgen TD (2001) Analysis of relative gene expression data using real-time quantitative PCR and the 2(-Delta Delta C(T)) Method. Methods 25:402–8 doi:10.1006/meth.2001.1262

Lozinsky OV, Lushchak OV, Kryshchuk NI, et al. (2013) S-nitrosoglutathione-induced toxicity in *Drosophila melanogaster*: Delayed pupation and induced mild oxidative/nitrosative stress in eclosed flies. Comp. Biochem. Physiol. A Mol. Integr. Physiol. 164:162–70 doi:10.1016/j.cbpa.2012.08.006

Lundberg JO, Weitzberg E (2022) Nitric oxide signaling in health and disease. Cell 185:2853–2878 doi:10.1016/j.cell.2022.06.010

Mense SM, Zhang L (2006) Heme: a versatile signaling molecule controlling the activities of diverse regulators ranging from transcription factors to MAP kinases. Cell Res. 16:681–92 doi:10.1038/sj.cr.7310086

Mercier A, Labbé S (2009) Both Php4 function and subcellular localization are regulated by iron via a multistep mechanism involving the glutaredoxin Grx4 and the exportin Crm1. J. Biol. Chem. 284:20249–20262 doi:10.1074/jbc.M109.009563

Mercier A, Watt S, Bahler J, Labbé S (2008) Key function for the CCAAT-binding factor Php4 to regulate gene expression in response to iron deficiency in fission yeast. Eukaryot. Cell 7:493–508 doi:10.1128/EC.00446-07

Mourer T, Brault A, Labbé S (2019) Heme acquisition by Shu1 requires Nbr1 and proteins of the ESCRT complex in *Schizosaccharomyces pombe*. Mol. Microbiol. 112:1499–1518 doi:10.1111/mmi.14374

Mourer T, Jacques JF, Brault A, Bisaillon M, Labbé S (2015) Shu1 is a cell-surface protein involved in iron acquisition from heme in *Schizosaccharomyces pombe*. J. Biol. Chem. 290:10176–90 doi:10.1074/jbc.M115.642058

Mourer T, Normant V, Labbé S (2017) Heme assimilation in *Schizosaccharomyces pombe* requires cell-surface-anchored protein Shu1 and vacuolar transporter Abc3. J. Biol. Chem. 292:4898–4912 doi:10.1074/jbc.M117.776807

Normant V, Brault A, Avino M, et al. (2021) Hemeprotein Tpx1 interacts with cell-surface heme transporter Str3 in *Schizosaccharomyces pombe*. Mol. Microbiol. 115:699–722 doi:10.1111/mmi.14638

Normant V, Mourer T, Labbé S (2018) The major facilitator transporter Str3 is required for low-affinity heme acquisition in *Schizosaccharomyces pombe*. J. Biol. Chem. 293:6349–6362 doi:10.1074/jbc.RA118.002132

Pelletier B, Trott A, Morano KA, Labbe S (2005) Functional characterization of the iron-regulatory transcription factor Fep1 from *Schizosaccharomyces pombe*. J. Biol. Chem. 280:25146–25161 doi:M502947200 [pii]; 10.1074/jbc.M502947200 [doi]

Peter C, Laliberté J, Beaudoin J, Labbé S (2008) Copper distributed by Atx1 is available to copper amine oxidase 1 in Schizosaccharomyces pombe. Eukaryot. Cell 7:1781–1794 doi:10.1128/EC.00230-08

Protacio RU, Mukiza TO, Davidson MK, Wahls WP (2022) Molecular mechanisms for environmentally induced and evolutionarily rapid redistribution (plasticity) of meiotic recombination. Genetics 220 doi:10.1093/genetics/iyab212

Reddi AR, Hamza I (2016) Heme Mobilization in Animals: A Metallolipid’s Journey. Acc Chem Res 49:1104–10 doi:10.1021/acs.accounts.5b00553

Rhee SG, Jeong W, Chang TS, Woo HA (2007) Sulfiredoxin, the cysteine sulfinic acid reductase specific to 2-Cys peroxiredoxin: its discovery, mechanism of action, and biological significance. Kidney Int. Suppl.(106):S3-8 doi:10.1038/sj.ki.5002380

Rhee SG, Woo HA (2011) Multiple functions of peroxiredoxins: peroxidases, sensors and regulators of the intracellular messenger H₂O₂, and protein chaperones. Antioxid Redox Signal 15:781–94 doi:10.1089/ars.2010.3393

Roy U, Yaish S, Weissman Z, et al. (2022) Ferric reductase-related proteins mediate fungal heme acquisition. eLife 11 doi:10.7554/eLife.80604

Rustici G, van Bakel H, Lackner DH, et al. (2007) Global transcriptional responses of fission and budding yeast to changes in copper and iron levels: a comparative study. Genome Biol. 8:R73 doi:10.1186/gb-2007-8-5-r73

Sabatinos SA, Forsburg SL (2010) Molecular genetics of *Schizosaccharomyces pombe*. Methods Enzymol. 470:759–795 doi:10.1016/S0076-6879(10)70032-X

Sarver A, DeRisi J (2005) Fzf1p regulates an inducible response to nitrosative stress in Saccharomyces cerevisiae. Mol. Biol. Cell 16:4781–91 doi:10.1091/mbc.e05-05-0436

Schmittgen TD, Livak KJ (2008) Analyzing real-time PCR data by the comparative C(T) method. Nature Protoc. 3:1101–8 doi:10.1038/nprot.2008.73

Sellam A, Tebbji F, Whiteway M, Nantel A (2012) A novel role for the transcription factor Cwt1p as a negative regulator of nitrosative stress in Candida albicans. PloS One 7:e43956 doi:10.1371/journal.pone.0043956

Severance S, Hamza I (2009) Trafficking of heme and porphyrins in metazoa. Chemical Rev. 109:4596–4616 doi:10.1021/cr9001116; 10.1021/cr9001116

Shakoury-Elizeh M, Tiedeman J, Rashford J, et al. (2004) Transcriptional remodeling in response to iron deprivation in Saccharomyces cerevisiae. Mol. Biol. Cell 15:1233–1243 doi:10.1091/mbc.E03-09-0642 [doi]; E03-09-0642 [pii]

Stojiljkovic I, Evavold BD, Kumar V (2001) Antimicrobial properties of porphyrins. Expert Opin. Investig. Drugs 10:309–20 doi:10.1517/13543784.10.2.309

Sung MK, Huh WK (2007) Bimolecular fluorescence complementation analysis system for in vivo detection of protein-protein interaction in *Saccharomyces cerevisiae*. Yeast 24:767–775 doi:10.1002/yea.1504

Sweeny EA, Singh AB, Chakravarti R, et al. (2018) Glyceraldehyde-3-phosphate dehydrogenase is a chaperone that allocates labile heme in cells. J. Biol. Chem. 293:14557–14568 doi:10.1074/jbc.RA118.004169

Swenson SA, Moore CM, Marcero JR, Medlock AE, Reddi AR, Khalimonchuk O (2020) From synthesis to utilization: The Ins and Outs of mitochondrial heme. Cells 9 doi:10.3390/cells9030579

Tillmann A, Gow NA, Brown AJ (2011) Nitric oxide and nitrosative stress tolerance in yeast. Biochem. Soc. Transactions 39:219–23 doi:10.1042/bst0390219

Tucker NP, Hicks MG, Clarke TA, et al. (2008) The transcriptional repressor protein NsrR senses nitric oxide directly via a [2Fe-2S] cluster. PloS One 3:e3623 doi:10.1371/journal.pone.0003623

Tucker NP, Le Brun NE, Dixon R, Hutchings MI (2010) There’s NO stopping NsrR, a global regulator of the bacterial NO stress response. Trends Microbiol. 18:149–56 doi:10.1016/j.tim.2009.12.009

Ullmann BD, Myers H, Chiranand W, et al. (2004) Inducible defense mechanism against nitric oxide in Candida albicans. Eukaryot. Cell 3:715–23 doi:10.1128/ec.3.3.715-723.2004

Vahsen T, Brault A, Mourer T, Labbé S (2023) A novel role of the fission yeast sulfiredoxin Srx1 in heme acquisition. Mol. Microbiol. 120:608-628 doi:10.1111/mmi.15146

Watanabe Y, Ishimori K, Uchida T (2017) Dual role of the active-center cysteine in human peroxiredoxin 1: Peroxidase activity and heme binding. Biochem Biophys Res Commun 483:930–935 doi:10.1016/j.bbrc.2017.01.034

